# A reconfigurable DNA memory architecture for hierarchical data management via programmable phase transitions

**DOI:** 10.64898/2026.01.17.700124

**Authors:** Jingyi Ye, Ao Liu, Fei Wang, Huan Zhang, Honglu Zhang, Chunhai Fan

## Abstract

DNA has emerged as a promising medium for the post-silicon era of information storage due to its ultrahigh density and longevity. However, current systems are bifurcated, with solid-state systems providing robust cold archival but lacking accessibility, while fluidic molecular computing systems offer dynamic processing but suffer from low density and instability. This mutual exclusivity has hindered the development of hierarchical memory, a standard in modern computing, within molecular storage systems. Here, we bridge this gap by engineering a reconfigurable DNA memory architecture driven by programmable liquid-liquid phase separation (LLPS). Our system leverages sequence-based encoding to achieve an ultrahigh storage density of 7×10^10^ GB/g, approaching the theoretical limits of DNA accessibility. In its fluidic hot state, DNA droplets enable rapid data loading (∼83.8% in 5 min) and function as an in-memory editing platform supporting versatile, addressable bit-level operations including selective erasure (∼65.1%) and high-efficiency rewriting and replacement (>99%) via programmable strand displacement. Importantly, to resolve the stability trade-off, we engineered a programmable phase transition whereby the triggered assembly of a rigid tetrahedral DNA framework (TDF) armor transforms liquid condensates into robust armored droplets. This cold state confers exceptional resistance to enzymatic and physical degradation, projecting multi-millennial data stability. By enabling reversible transitions between a editable, high-density computing mode and a stabilized archival mode, this work establishes the architectural foundation for scalable molecular information storage capable of hierarchical data management.

## Introduction

The exponential growth of digital data is pushing conventional silicon-based memory technologies toward their physical and economic limits^1, 2^. As magnetic and optical storage media approach their scaling boundaries, DNA has emerged as a promising alternative, offering information densities orders of magnitude higher than electronic storage, together with exceptional energy efficiency and millennial-scale durability^3–5^. However, translating DNA from a biological archive into a practical information substrate faces a fundamental architectural challenge. Modern computing systems do not treat all data uniformly. Instead, they rely on a hierarchical memory architecture that distinguish between hot data, which are frequently accessed, modified and processed, and cold data consisting of archival records requiring long-term preservation with minimal intervention^6–9^. Reproducing this functional hierarchy in molecular systems remains a central obstacle to advancing DNA storage beyond static archiving.

To date, progress in DNA-based storage technologies has largely followed two parallel yet distinct directions^4^. One path emphasizes cold archival preservation through embedding DNA within protective matrices such as silica beads^10^, hydrogels^11^, liquid crystal^12^, or metal-organic frameworks (MOFs)^13^ to shield information from entropy. While these approaches provide robust protection against environmental degradation, they are incompatible with active data management; accessing or modifying data often requires chemical dissolution or mechanical separation that destroys the storage architecture^14, 15^. The other developmental path focuses on dynamic molecular operations, utilizing exposed DNA strands in solution to perform logic and computation^16–21^. Recent studies have shown the potential of DNA condensates for information encryption and logic operations^22^. However, a critical bottleneck remains in bridging these dynamic capabilities with the high-density requirements of archival storage. Current fluidic systems often rely on spatial or fluorescence-based encoding approaches that sacrifice information density and long-term protection compared to sequence-based archival systems. Consequently, the field currently lacks a mechanism to seamlessly toggle a single system between a reactive, editable state for high-density computation and a rigid, inert state for long-term archiving, a gap that precludes the development of molecular hierarchical memory^4^.

Here, we address this challenge by developing a reconfigurable DNA memory architecture that unifies dynamic hot-data editing with stable cold-data archival within a single platform (Figure 1). Rather than relying on static containment, our approach leverages liquid-liquid phase separation (LLPS) of DNA to create a programmable state-switching storage medium. In its hot state, the system forms fluid DNA droplets that enable rapid data loading (∼83.8% within 5 min), and provide molecular access for in-situ file operations. Using sequence-based encoding, this platform achieves an ultrahigh storage density of approximately 7×10^10^ GB/g, approaching the theoretical limit for accessible DNA, while supporting addressable erasure (∼65.1% efficiency) and high-efficiency rewriting and replacement (>99%) via programmable strand displacement. Importantly, to complete the data life-cycle, we introduce a programmable state lock. By engineering self-assembling tetrahedral DNA frameworks (TDFs) to interfacially assemble at the droplet surface, we trigger an on-demand transition from the fluid hot state to an armored state for cold archiving. This molecular armor forms a physical barrier that protects the encoded DNA against enzymatic degradation and physical stress, yielding projected multi-millennial stability comparable to solid-state encapsulation. The ability to reversibly transition between an editable and a protected state establishes a reconfigurable DNA memory architecture based on hierarchical data management, providing a foundation for practical and scalable molecular data storage.

**Figure 1.**
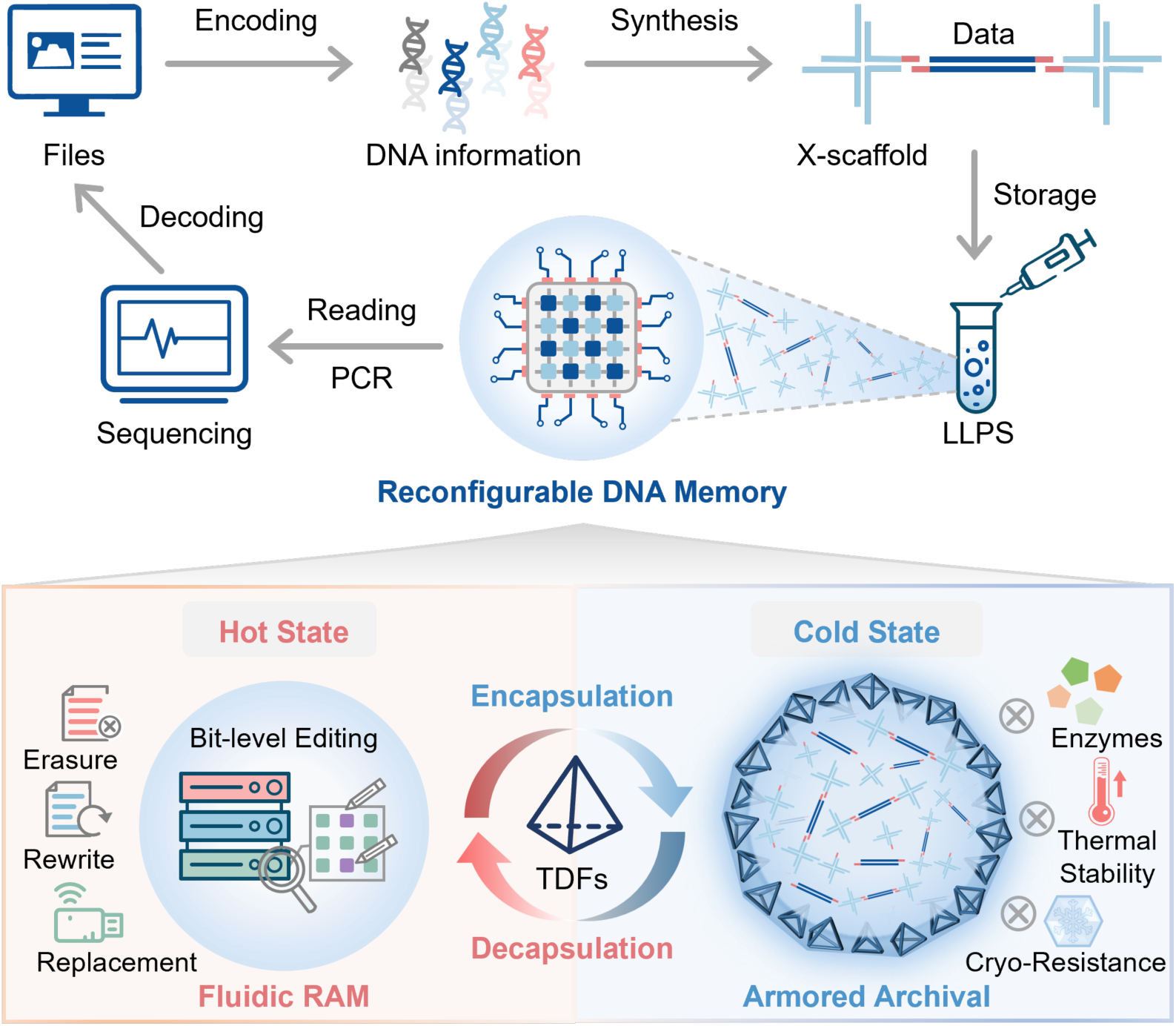
A phase-transitioning DNA memory architecture. Top: Workflow converting digital files into sequence-based condensate droplets via LLPS. Bottom: Programmable transition between RAM Mode (Left), where fluidic droplets enable rapid in-memory editing (erasure, rewriting, replacement), and Archive Mode (Right), where TDF encapsulation forms a robust armor against enzymes, heat, and cryopreservation. Center: The cycle depicts the reversible toggling between the editable fluidic state and the protected archival state.

## Results

### Tunable condensate fluidity governs hot data loading kinetics

To engineer an effective hot state capable of rapid data exchange, we first established a DNA condensate system driven by multivalent interactions between X-scaffolds and data strands. While liquid-liquid phase separation (LLPS) drives droplet formation, the physical state of the condensate ranging from liquid to gel-like, is a critical determinant of data management efficiency^23, 24^. To investigate the relationship between scaffold design and loading kinetics, we systematically modulated the sticky-end interactions governing network assembly through varying both length (4-nt vs. 6-nt) and sequence symmetry (palindromic, P vs. non-palindromic, NP) (Figure 2a). We found that thermodynamic stability is a essential for efficient data loading. The 4-nt designs, regardless of symmetry, exhibited minimal payload uptake due to weak intermolecular affinity (Figure 2b-2c and Figure S1-S4). In contrast, 6-nt designs formed data-enriched condensates but revealed a sharp divergence in loading kinetics based on sequence symmetry. The non-palindromic NP-6 system achieved rapid and high-capacity loading, reaching near-saturation within 15 minutes. In stark contrast, the palindromic P-6 system exhibited significantly slower kinetics, requiring 60 minutes to reach a comparable payload level.

**Figure 2.**
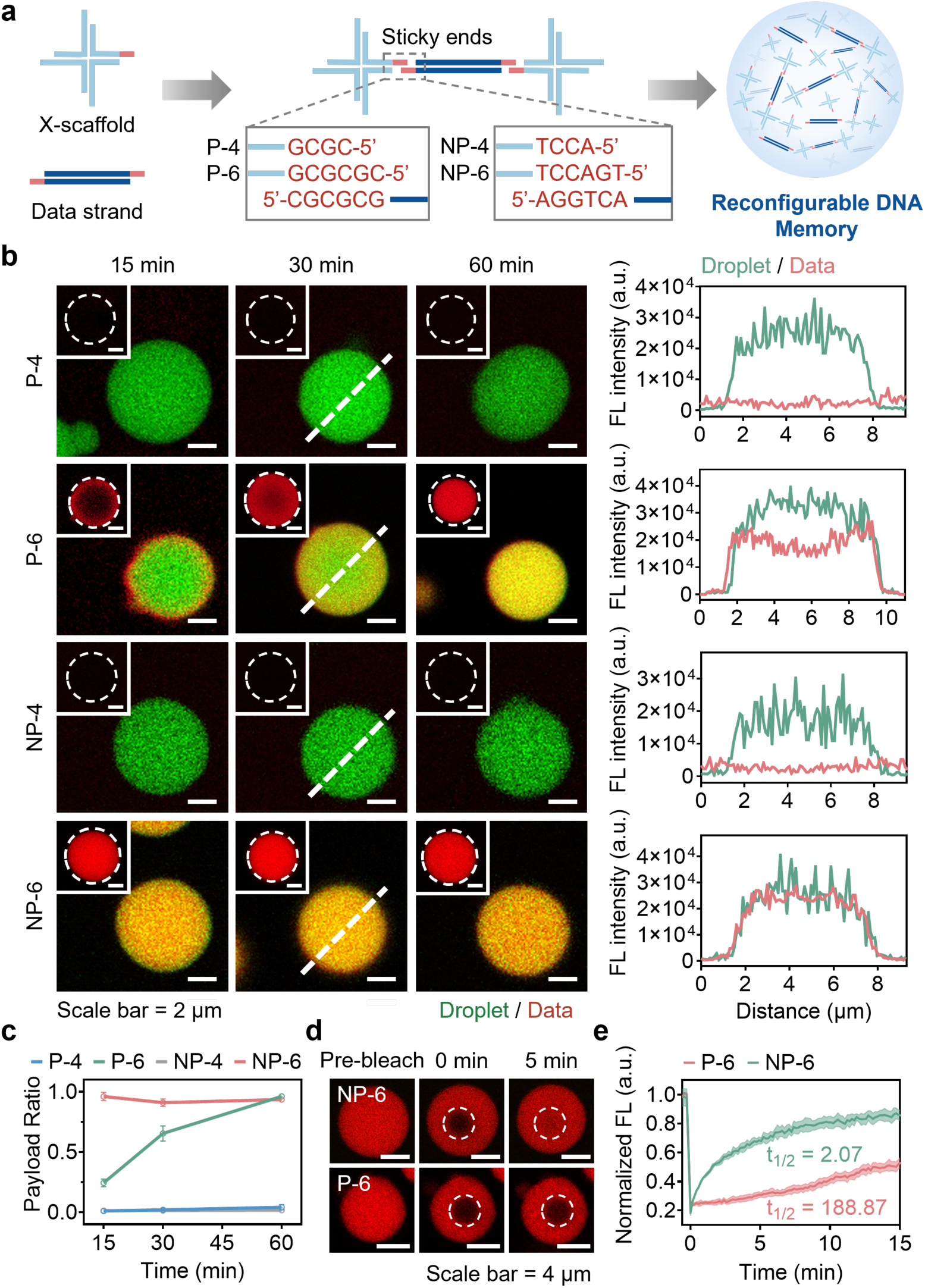
Regulation of condensate fluidity and data loading kinetics via sticky-end topology. **a,** Schematic of the sticky ends designs (P-4, P-6, NP-4, NP-6) used to modulate the physical state and loading efficiency of the DNA memory. Designs vary in overhang length (4-nt vs. 6-nt) and sequence symmetry (Palindromic, P vs. Non-Palindromic, NP). **b,** Temporal fluorescence imaging (left) and corresponding intensity profiles (right) showing the distinct data (50 bp dsDNA) loading behaviors across the four sticky-end designs. While 4-nt designs show negligible uptake, 6-nt designs exhibit symmetry-dependent kinetics. The white dashed circle indicates the region of interest (ROI) for intensity analysis at the 30-min time point. **c**, Quantitative kinetics of data internalization plotted as the payload ratio (Data/Droplet) over time. **d-e,** Probing internal diffusivity via FRAP analysis. **d**, Representative pre- and post-bleach images comparing the recovery dynamics of P-6 and NP-6 droplets. **e**, Normalized fluorescence recovery curves reveal that NP-6 droplets exist in a highly fluid, liquid-like state, whereas P-6 droplets form kinetically trapped gels. All experiments utilized 500 nM data strands (Cy5, red) and 5 µM X-scaffolds (FAM, green). Error bars represent SD (n = 3).

To clarify the mechanistic basis of this kinetic difference, we examined the internal diffusivity of the condensates using fluorescence recovery after photobleaching (FRAP)^25^. The NP-6 droplets displayed rapid and complete fluorescence recovery (t_1/2_ = 2.07 min), confirming a highly fluid, liquid-like microenvironment critical for efficient molecular transport (Figure 2d-2e). However, P-6 droplets behaved like kinetically trapped gels (t_1/2_ = 188.87 min), showing dynamics approximately 91 times slower than the NP-6 system. This gelation resulted from the strong self-complementary interactions between palindromic sticky ends, which create a rigid scaffold network that severely restricts the diffusion and infiltration of data strands. These findings highlight a fundamental design principle: internal fluidity, regulated by the topology of the scaffold network, governs data-loading kinetics. Maintaining a liquid-like phase is therefore not just a physical property but a functional prerequisite for high-speed hot data memory. Based on these insights, the NP-6 design, which optimally combines the high binding affinity with the necessary fluidity of non-palindromic interactions, was selected as the platform for all subsequent operations.

### Ultrahigh density storage with rapid writing and direct accessibility

Having engineered the optimal fluidic platform, we next characterized its core performance as a storage architecture, focusing on the critical triad of writing speed, storage density and data accessibility. First, regarding writing kinetics, the fluidic nature of the NP-6 droplets enabled the rapid internalization of data strands across diverse lengths (50, 75, and 100 bp), with loading reaching saturation within approximately 15 minutes (Figure 3a). This behavior indicates broad compatibility with variable payload sizes and efficient uptake across sequence lengths relevant to practical data encoding. Notably, kinetic analysis of 100 bp dsDNA revealed an particularly rapid absorption rate, achieving approximately 83.8% of maximum loading capacity within just 5 min (Figure 3b and Figure S5). These results highlight the suitability of the condensate-based system for time-sensitive hot data management, compared with solid-state preservation approaches that typically require slower and multistep encapsulation procedures.

**Figure 3.**
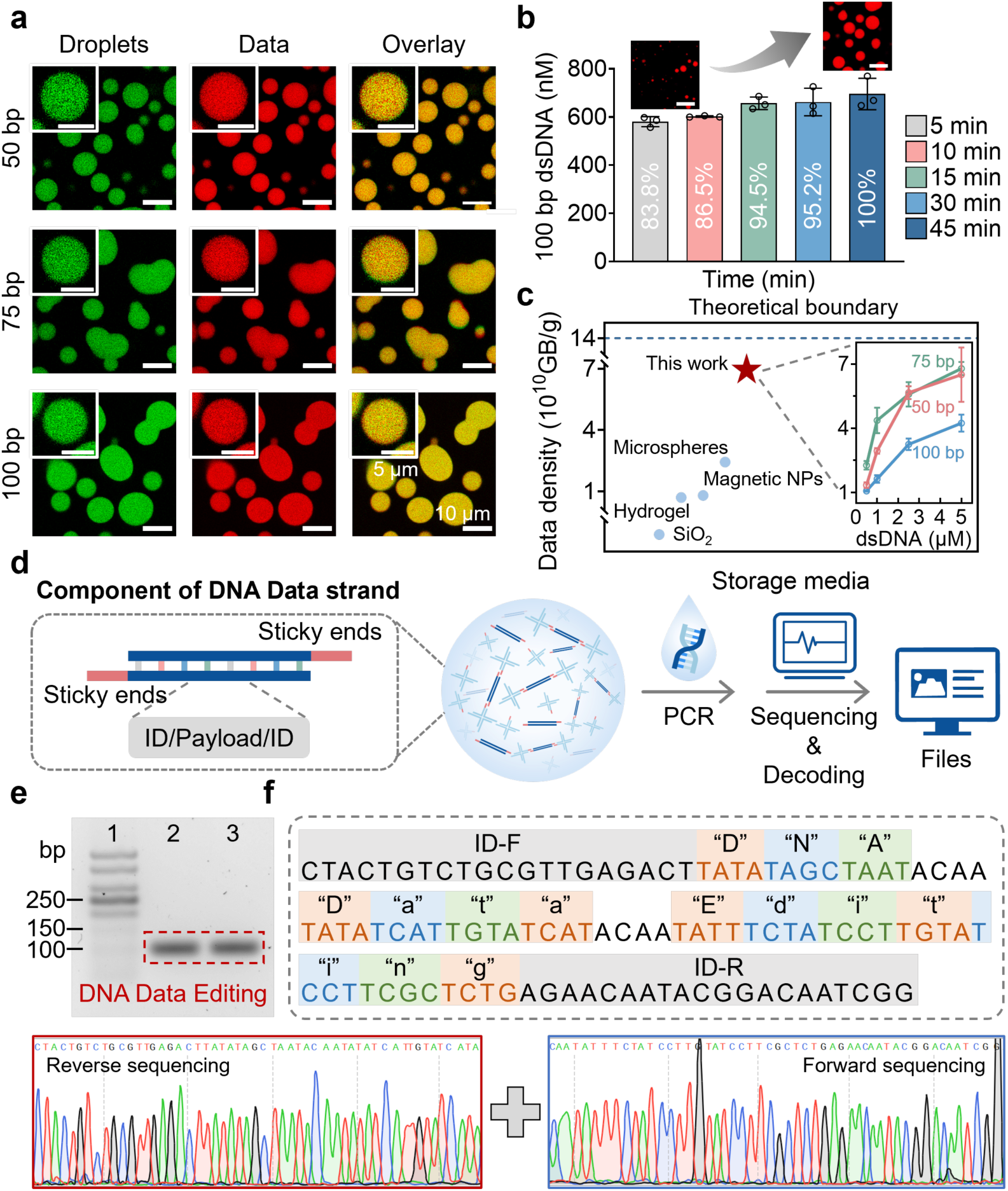
Ultrahigh density storage and direct accessibility of the fluidic hot phase. **a,** Confocal fluorescence images demonstrating the universal loading capability of NP-6 droplets for data strands of varying lengths (50, 75, and 100 bp). **b,** Fast-writing kinetics. Top: Representative images of 100 bp dsDNA (red) internalized in 5 min to 45 min. Bottom: Quantitative analysis shows rapid uptake, reaching ∼83.8% of capacity within 5 minutes. Scale bar, 10 μm. **c**, Comparison of data density across different media for DNA data storage, including SiO_2_^18^, TRFG hydrogels^11^, multilayer magnetic nanoparticles^30^, photonic microspheres^13^ and this work. The theoretical boundary of this work is ∼14×10^10^ GB/g (details in Supplementary Methods). Quantification of storage density as a function of data strand concentration and length (inset). The system achieves a peak physical density of ∼7×10^10^ GB/g (with 75 bp strands at 5 μM), validating the efficiency of sequence-based encoding. **d**, Schematic of the direct readout workflow and data strand architecture, comprising a central information payload flanked by universal primer-binding sites (ID domains) for non-destructive retrieval. **e,** Gel electrophoresis verification of the PCR product amplified directly from the liquid droplets, confirming the successful retrieval of the file “DNA Data Editing”. **f,** Verification of data fidelity via bi-directional Sanger sequencing and decoding of the amplicons from (**e**).

Crucially, we sought to resolve the trade-off between operational capability and storage density. While recent storage systems have relied on spatial-fluorescence^22^ or structure encoding strategies^26, 27^, we used a sequence-based encoding strategy to fully exploit the information capacity of the DNA molecule, in which each nucleotide encodes two bits of information^1, 28^. By quantifying payload uptake under varying strand concentrations, we observed that storage density is governed by a balance between strand availability and steric hindrance of DNA cargo. At high concentration of data strands (5 μM), the system achieved a peak density of ∼7×10^10^ GB/g using 75 bp payloads (Figure 3c). While the density for 100 bp strands was slightly lower (∼4.2×10^10^ GB/g) due to increased steric hindrance within the condensate network. These values place our fluidic memory among the highest-density DNA storage platforms reported to date (Figure 3c and Table S1). This achievement demonstrates that the hot state can approach the theoretical limits of molecular density while retaining the fluidic accessibility required for dynamic management.

Beyond high-density storage, a practical memory architecture must support efficient and non-destructive data retrieval. Unlike solid encapsulation strategies (e.g., DNA embedded in silica or MOFs) that typically require chemical dissolution or physical destruction to access stored information^14, 15, 29^, our DNA memory remains chemically accessible in its fluid state. To demonstrate this capability, we designed data strands containing a central payload domain flanked by primer-binding sites (ID domains, Figure 3d). After storing the ASCII-encoded message “DNA Data Editing”, we directly amplified the target sequences from the droplet suspension using PCR without any prior release or purification steps. The retrieved information was verified by agarose gel electrophoresis (Figure 3e) and Sanger sequencing (Figure 3f), confirming accurate and high-fidelity recovery. Thus, these results demonstrate that the system integrates ultrahigh storage density with direct and non-destructive access, providing a robust framework for precise hot data operations within a fluid molecular environment.

### Programmable in-memory logic and erasable rewriting in the fluidic phase

A defining feature of hot data management is the capacity for frequent random-access updates, a functionality that is routine in silicon random-access memory (RAM) but has remained challenging in molecular storage^32–34^. Having validated the fluidic accessibility of the NP-6 droplets, we next configured the system to operate as a dynamic in-memory editing platform. To enable orthogonal addressability, we constructed a model file library containing two distinct data populations, Data A and Data B. While sharing a common sticky-end domain (m*) to allow uniform uptake into the droplets, each file incorporates a unique toehold domain (n* and f*) that serves as a specific address for sequence-specific operations (Figure 4a).

**Figure 4.**
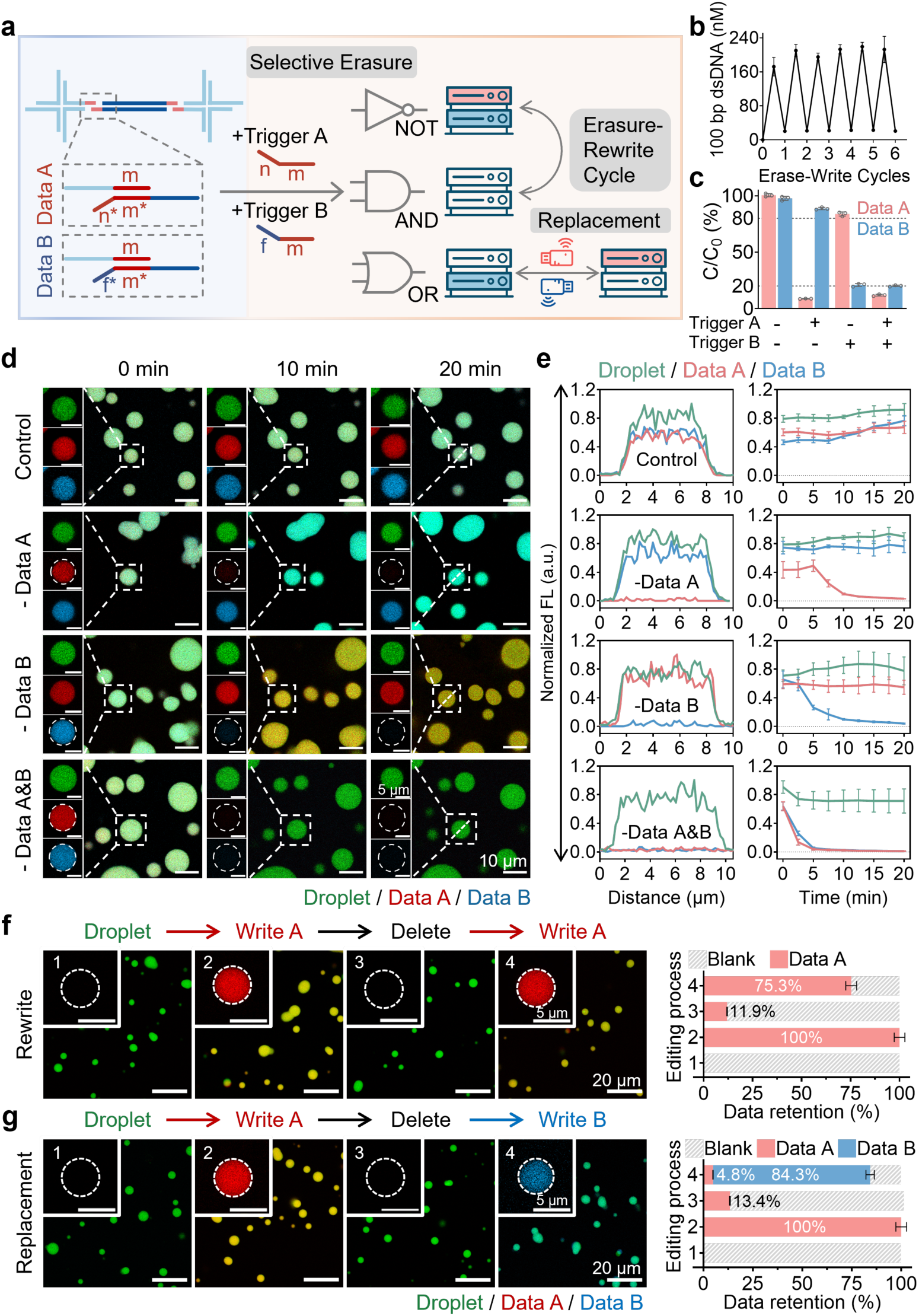
Programmable in-memory logic and erasable rewriting. **a**, Schematic of the addressable operation logic. Data A and Data B share a storage domain (m*) but possess orthogonal toeholds (n* and f*) to enable selective strand displacement (erasure) and independent rewriting or replacement. **b**, Cyclic endurance test showing the concentration of 100 bp dsDNA over six consecutive erase-rewrite cycles, demonstrating the robust reusability of the fluidic phase. **c**, Logic-gated erasure efficiency. Quantitative PCR analysis of Data A (red) and Data B (blue) retention following specific trigger inputs (NOT, OR, AND gates), confirming high orthogonality. **d**, Time-lapse confocal microscopy visualizing the selective removal of specific data files (Data A, Data B, or both) from the droplets within minutes. **e**, Normalized fluorescence intensity profiles (left) and temporal kinetics (right) extracted from (**d**), highlighting the rapid response of the system. **f-g**, Demonstration of complete data management workflows. Representative images (left) and quantification (right) of data retention during a rewriting cycle (**f**, Blank-Written Data A-Erased-Rewritten Data A) and a replacement cycle (**g**, Blank-Written Data A-Erased-Replaced Data B). The system achieves high fidelity in both restoring original data and swapping file content. Error bars represent SD (n = 3).

We first evaluated the system’s endurance under repeated editing cycles, a key requirement for dynamic memory. We achieved data selective erasure via toehold-mediated strand displacement, in which trigger strands selectively recognize and release the target data strands from the droplets. The high internal fluidity of the NP-6 phase ensures that these reactions proceed without diffusion limitations, avoiding the kinetic trapping commonly encountered in more rigid DNA materials^35, 36^. Across six consecutive erase-rewrite cycles, the overall data concentration remained stable, with no observable degradation of droplet structure or phase behavior (Figure 4b). The minor fluctuations in signal intensity were attributable to wash-step losses rather than intrinsic system fatigue, indicating that the platform supports repeated data updating with minimal hysteresis.

Building on this capability, we implemented programmable logic operations directly within the droplets using sequence-specific trigger strands. By selectively addressing Data A, Data B, or both, we realized a set of molecular logic gates including NOT (Control), OR (Delete A or B) and AND (Delete A & B) operations, to execute precise data manipulation. The sequence-based strand displacement approach offers high-dimensional programmability for our whole DNA-based storage platform through orthogonal toehold designs. Quantitative analysis showed efficient and selective erasure, with removal efficiencies of approximately 91.3% for Data A and 79.1% for Data B, accompanied by limited off-target effects of around 13.8% (Figure 4c). The difference in erasure efficiency likely reflects variations in toehold binding kinetics^37^. Importantly, this modest level of crosstalk did not compromise functionality, as it was readily corrected during subsequent rewriting steps performed at higher strand concentrations. Confocal microscopy visualized this process in real-time, showing the rapid and selective removal of fluorescent data signals within minutes (Figure 4d-4e and Figure S6-S9). Notably, the dual-target erasure (AND gate) kinetics (∼5 min) proceeded faster than single-target erasure (OR gate, ∼10 min), suggesting cooperative acceleration of strand displacement within the crowded yet fluid microenvironment. This kinetic advantage highlights the suitability of the liquid condensate phase for rapid and parallel processing.

Finally, we combined these operations to demonstrate a complete write-erase-rewrite/replace workflow, simulating the full lifecycle of active data management. The memory state was sequentially transitioned from an initial blank state to written, erased, and subsequently rewritten or replaced states (Figure 4f-4g). The system exhibited high operational fidelity, with rewriting restoring ∼75.3% of the original signal and replacement achieving ∼84.3% loading efficiency. These results confirm that our DNA memory functions not only as a high-density storage medium, but as a responsive molecular system capable of supporting the precise, repeatable and kinetically efficient operations required for scalable hot data management.

### Scalable, addressable bit-level editing in complex data environments

A major challenge in advancing molecular storage is achieving fine-grained data manipulation, specifically the ability to perform addressable, bit-level updates within a complex library without disrupting the surrounding dataset^4^. Such localized editing is critical for practical file management, as it offers enhanced operational efficiency compared to the costly global rewriting required by current static storage media. However, many existing dynamic systems lack this level of granularity, requiring physical separation or bulk reprocessing to modify specific data owing to intrinsic encoding limitations^22, 27^. In contrast, our fluidic architecture combines the high programmability of DNA strand displacement with an open, diffusion-enabled network. This design creates a a memory environment analogous to electronic RAM and allows targeted modification of specific data strands while preserving the integrity of the remaining library.

To validate fine-grained addressability, we first engineered a proof-of-concept text file encoded as a small library of six distinct DNA strands (135 bytes total, Figure 5a). Each strand was designed with a unique address index and orthogonal toehold domains (Figure S10), enabling the specific targeting required for bit-level operations (Supplementary Methods and Supplementary Note 1). Illumina sequencing confirmed that all strands maintained read counts well above the decoding threshold, with minor variations in abundance attributable to GC content (Figure 5b). To assess operational precision, we performed a targeted delete-rewrite-replace workflow on two selected strands Txt_2 and Txt_6. Following toehold-mediated deletion, the retention of target strands (Txt_2a and Txt_6a) dropped significantly by approximately 65.9%, while non-target strands remained unchanged, confirming the high specificity of the erasure operation. A subsequent rewrite operation successfully restored their abundance to ∼74.5% of the original level (Figure 5c and Figure S11), demonstrating reversible and controlled editing. Beyond restoration, we executed a “digital mutation” by introducing modified strands (Txt_b) to replace the original ones (Txt_a). Sequencing analysis revealed a clear population shift in which the replacement strands dominated the read pool, while the original strands were reduced to low residual levels (Figure 5d). Our decoding algorithm successfully filtered this residual noise to identify the updated information (Supplementary Methods and Supplementary Note 2). As a result, this high-fidelity process enabled the precise semantic transformation of the stored text from “a grain of sand” to “a droplet of DNA” and “an hour” to “a moment” (Figure 5e), verifying the the system supports precise, context-aware bit-level editing rather than simple file-level rewriting.

**Figure 5.**
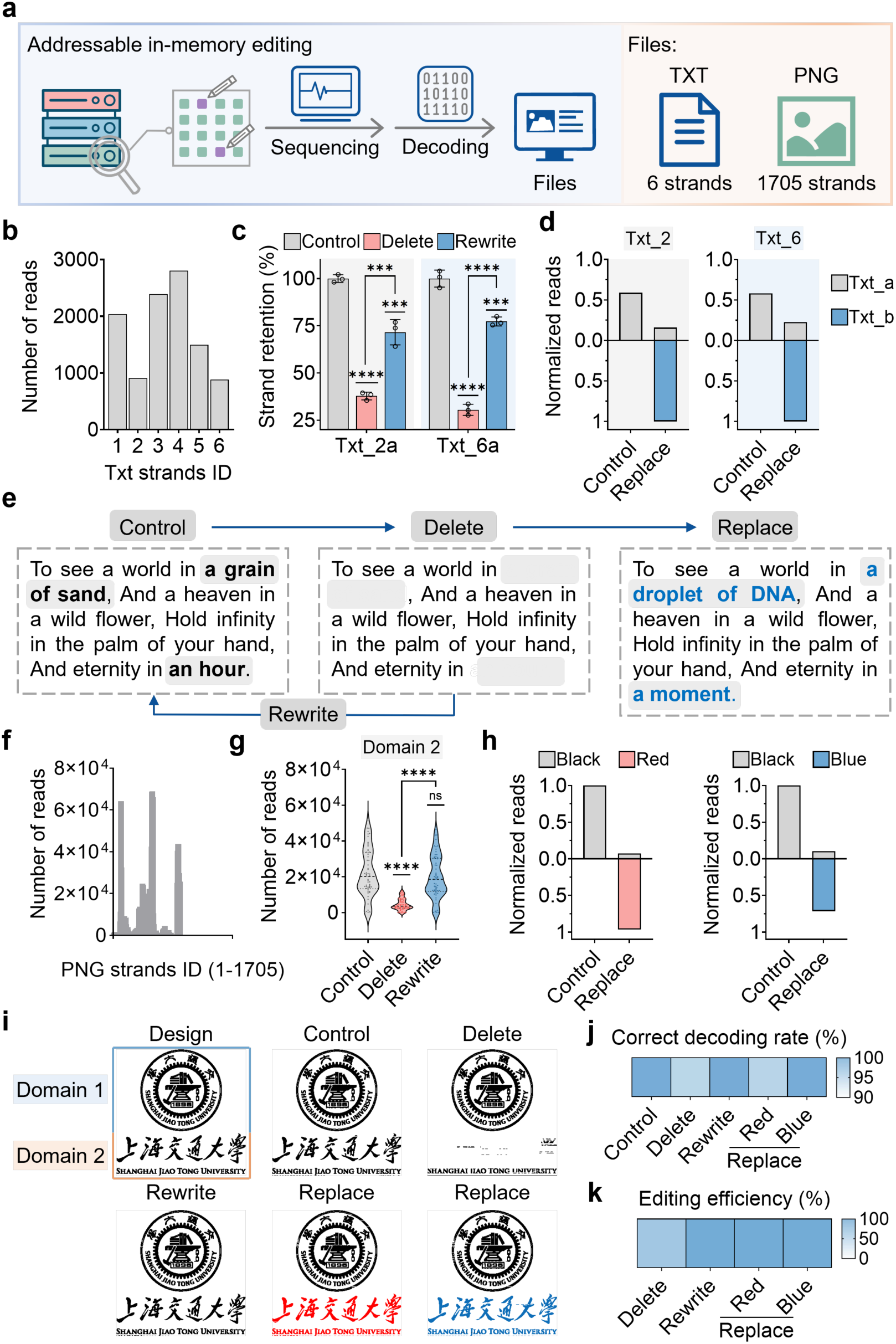
Scalable and precise bit-level editing in complex data environments. **a**, Workflow for addressable bit-level operations. The system was validated across two scales of complexity: a simple 6-strand TXT file and a complex 1,705-strand PNG file. **b**, Read distribution profiling of the initial TXT library via Illumina sequencing. **c**, Targeted manipulation efficiency. Retention ratios of specific strands (Txt_2a, Txt_6a) following selective deletion and rewriting operations demonstrate high responsiveness. **d**, Replacement efficiency. Normalized sequencing reads show the dominant emergence of replacement strands (Txt_b, blue) over original strands (Txt_a, gray). **e**, Semantic editing demonstration. Successful decoding of the TXT file reveals the precise transformation of text content. **f**, Read distribution of the complex 1,705-strand PNG library, serving as a high-noise testbed. **g**, Domain-specific editing. Significant read count shifts in the editable Domain (Domain 2) confirm targeted operations within the large pool. **h**, Color-change operation. Normalized reads confirm the successful replacement of “Black” coding strands with “Red” and “Blue” strands. **i**, Visual decoding results. The university badge is successfully modified from its original design to updated color variants. **j-k**, Performance metrics. The system achieves a ∼97.7% correct decoding rate (**j**) and >99% rewriting/replacement and ∼65.1% selective erasure efficiency (**k**) in the complex PNG environment. Error bars represent SD (n = 3). Statistical significance determined by one‐way ANOVA (***p < 0.001, ****p < 0.0001; ns, p > 0.05).

To further challenge the system’s scalability and specificity in a high-complexity environment, we extended the approach to a complex PNG image encoded by 1,705 distinct DNA strands representing a university badge. This high-complexity library introduces significantly higher background noise and therefore serves as a stress test for hybridization specificity. The dataset was partitioned into a static background region (Domain 1) and an editable foreground region (Domain 2). Initial sequencing confirmed adequate coverage across the full 1,705-strand pool (Figure 5f), establishing a robust baseline for complex file storage. Targeted editing was then performed exclusively on the foreground region to alter the color encoding of the badge. Following deletion and rewriting, we observed a significant and targeted shift in the read counts of Domain 2 strands compared to the original control group, confirming that the hot operations remain orthogonal even in a crowded molecular environment (Figure 5g). Furthermore, during the replacement operation (switching from black to red/blue coding strands), the normalized read counts for the new color strands were significantly higher than those for the original strands (Figure 5h), yielding a high signal-to-noise ratio critical for accurate decoding. This selective enrichment enabled correct reconstruction of the modified badge image (Figure 5i). Quantitative analysis across these large-scale editing experiments showed an average decoding accuracy of approximately 97.7%, with selective erasure efficiencies of around 65.1% and rewriting and replacement fidelities approaching 99.7% (Figure 5j-5k). These results demonstrate that the our DNA memory supports precise and scalable bit-level editing even within complex and noisy data environments, validating its potential for dynamic hot data management at practical scales.

### Programmable phase transition for reversible and robust archival

A hierarchical memory architecture must support not only dynamic editing but also the reliable conversion of active data into a stable archival state. To meet this requirement, we developed a programmable phase transition that reversibly switches the memory between a fluidic editable mode and a rigid archival mode. Unlike conventional core-shell designs that permanently entrap DNA inside silica or polymer matrices, our approach uses tetrahedral DNA frameworks (TDFs) to construct a reconfigurable protective armor around the data-enriched droplets (Figure 6a). The transition is driven by a triggered self-assembly process. Bridge strands recruit TDFs onto the droplet interface, where palindromic vertex overhangs facilitate lateral cross-linking to form a mechanically rigid armor. Importantly, this interfacial phase transition is fully reversible. The addition of specific release strands initiates a competitive toehold-mediated displacement that detaches the TDF armor, restoring the droplets to their fluid state. Confocal microscopy directly visualized this reversible lock-unlock cycle, with TDFs confined to the droplet periphery in the archival state (Figure 6b and Figure S12) and completely removed upon reversion to the hot state (Figure 6c and Figure S13). This reversibility enable data to be protected for archival and subsequently reactivated for on-demand editing.

**Figure 6.**
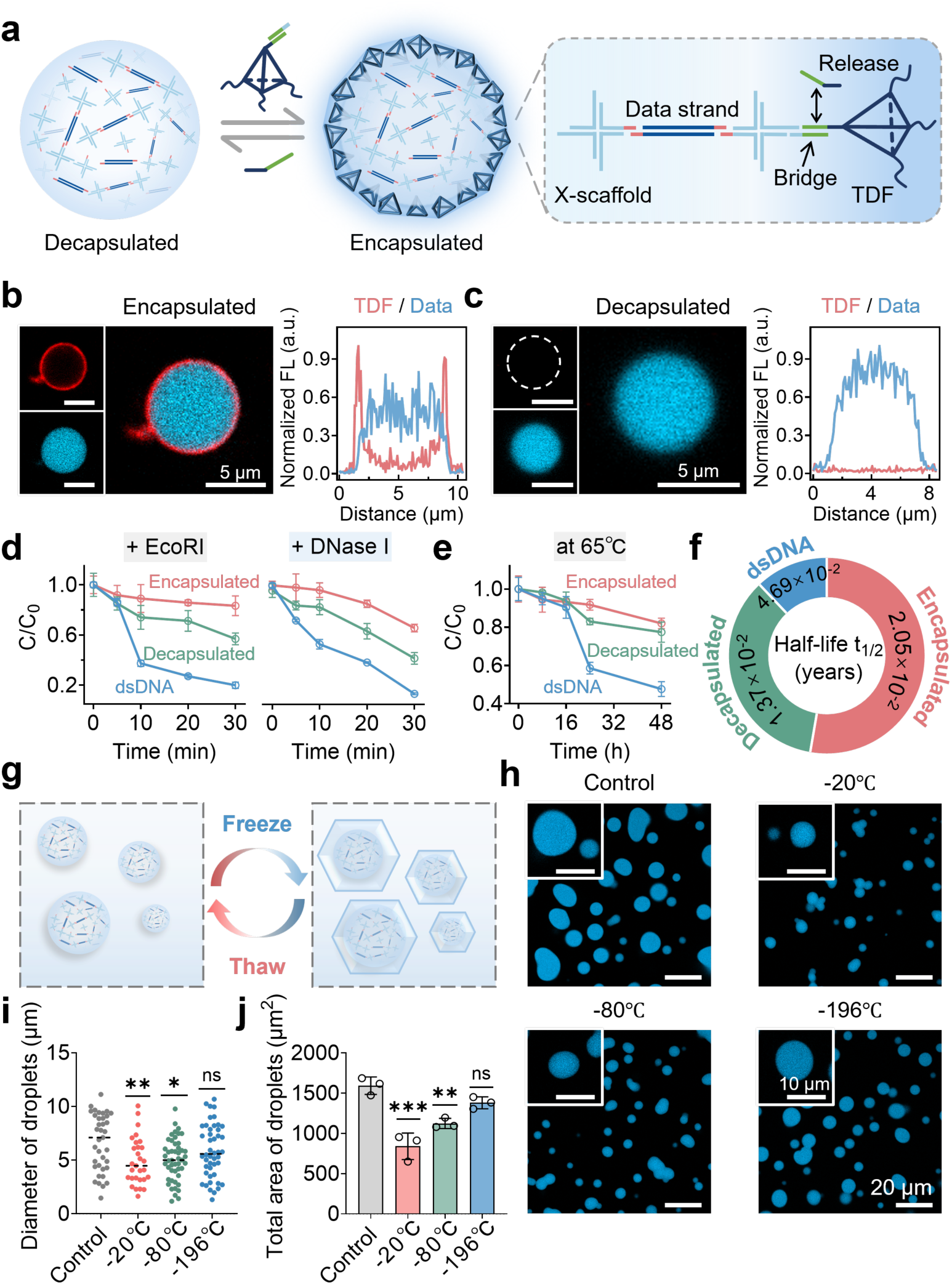
Programmable phase transition for reversible and robust cold archival storage. **a,** Schematic of the reversible encapsulation mechanism. Triggered assembly of TDFs forms a rigid armor (cold State) for protection, while release strands detach the armor via strand displacement to restore fluidity (hot State). **b-c**, Visual confirmation of the phase transition. Confocal images (left) and intensity profiles (right) showing the precise surface localization of TDFs (Cy5, red) upon encapsulation (**b**) and their complete removal upon decapsulation (**c**). Scale bars, 5 µm. **d**, Enzymatic resistance profiling. Degradation kinetics of data strands in free solution (blue), decapsulated droplets (green), and encapsulated droplets (red) upon exposure to EcoRI (left) and DNase I (right). **e**, Accelerated aging kinetics. Decay of data concentration over 48 h at 65°C, highlighting the stabilizing effect of the TDF armor. **f**, Projected longevity. Half-life (t_1/2_) estimations extrapolated from (**e**), predicting multi-millennial stability for the encapsulated state. **g**, Workflow for cryopreservation stress testing across different thermal regimes. **h**, Post-thaw structural integrity. Fluorescence images of data strands (blue) following retrieval from −20°C, −80°C, and −196°C storage. Liquid nitrogen storage (−196°C) preserves morphology best by preventing ice crystal formation. **i-j**, Statistical analysis of droplet diameter (**i**) and total area (**j**) post-thaw, confirming that vitrification at −196°C causes negligible physical damage. Error bars represent SD (n = 3). Statistical significance determined by one‐way ANOVA (*p < 0.1, **p < 0.01, ***p < 0.001; ns, p > 0.05).

In addition to reversibility, this architecture introduces physical discretization, which is essential for scalable storage. Fluid droplets in the hot state readily fuse, leading to loss of addressability through macroscopic phase separation. Encapsulation with a TDF armor suppresses this coalescence, preserving the droplets as stable, monodisperse units even upon contact (Figure S14). This design biomimics a digital folder system, in which files are maintained as distinct logical units. Although sequence-based indexing allows data retrieval in principle, physical compartmentalization prevents aggregation and sedimentation, ensuring long-term manageability of the archival medium. We next quantified the protective performance of the TDF armor under stringent stress conditions. When exposed to restriction enzymes (EcoRI) or non-specific nucleases (DNase I), encapsulated data showed markedly enhanced resistance compared to naked DNA or unshielded droplets, confirming that the armor acts as an effective semi-permeable barrier (Figure 6d). Accelerated aging experiments at 65 °C further demonstrated strong suppression of hydrolytic decay. The measured decay rate was 1.07×10^-6^ s^-1^ (corresponding to a half-life of 0.0205 years) at 65°C and an extrapolated half-life of approximately 4,000 years at 4 °C (Figure 6e-6f, Table S2). Based on Arrhenius extrapolation models^38, 39^, this stability extends to approximately 1.4 million years under conditions mimicking the Global Seed Vault (−18°C), rivaling the longevity of fossilized DNA (Supplementary Methods).

Finally, we tested whether the encapsulated system could withstand extreme bulk phase transitions associated with cryopreservation. We subjected data-loaded droplets to freezing at −20°C, −80°C, and −196°C (Figure 6g). Our results revealed that slow freezing (−20°C and −80°C) caused structural damage due to ice crystal formation^40^, while rapid vitrification in liquid nitrogen (−196°C) fully preserved droplet integrity. After thawing, vitrified droplets showed complete morphological restoration with no significant changes in size or surface area relative to unfrozen controls (Figure 6h-6j). Together, these results demonstrate a programmable interface strategy that converts a dynamic DNA memory into a robust archival vault. The self-assembled TDF armor provides multi-millennial protection against enzymatic, thermal, and physical degradation while remaining fully reversible. By uniting fluidic accessibility with rigid preservation in a single platform, this system completes the molecular data life cycle and offers a molecular implementation of hierarchical data management.

## Conclusions

In summary, we have established a reconfigurable DNA memory architecture that resolves the architectural dichotomy between dynamic data processing and robust archival preservation. By seamlessly bridging fluidic accessibility and switchable structural protection in a single system, our platform moves beyond the intrinsic limitations of both static, write-once media and exposed solution-phase pools. At the core of this advance is a programmable mode reconfiguration enabled by LLPS and DNA nanotechnology, alowing the memory to transition on demand between a kinetically active hot state and a stable cold state.

Our architecture represents distinct advances over existing DNA storage paradigms, which often lack the systemic integration required for hierarchical memory (Table S3). While solution-based systems offer accessibility^16, 17^, they leave information chemically exposed, lacking the structural protection necessary for long-term archiving. In contrast, solid or material-based carriers^21^ improve stability but typically rely on irreversible encapsulation or coarse release release mechanisms that restrict editing precision. Recent condensate-based systems have begun to explore core-shell concepts^22^, they have largely focused on spatial encoding for logic operations, with limited storage density and archival robustness. Our work advances this domain by combing sequence-based encoding and programmable TDF encapsulation, thereby uniting ultrahigh information density, bit-level editability, and long-term protection within a single architecture.

A key advance of this system is the reversibility of the transition between hot and cold states. Rather than functioning as a permanent molecular fossil, the encapsulated state acts as a programmable lock that preserves information over extended timescales while retaining the option for future access. Data can be stabilized against environmental stress and later returned to a fluid, editable form when updates are required. This reversible control closes the data lifecycle, transforming DNA storage from a passive archive into an adaptable memory that supports both preservation and evolution of information.

Looking forward, practical implementation must address challenges such as synthesis costs, error control and large-scale operation^41–44^. Nevertheless, the modular and fluidic design of this architecture makes it well compatible with emerging technologies like enzymatic DNA writing and microfluidic automation^45–47^. Importantly, encapsulation introduces physical discretization as a structural feature for information handling at the level of individual files. Unlike bulk solutions, TDF-encapsulated units serve as addressable, non-coalescing compartments, mimicking the file folder organization of digital storage. When combined with automated tools such as microfluidic routing or fluorescence-activated cell sorting (FACS), this system enables physically addressable molecular memory, where data can be selectively located, retrieved, and processed without global manipulation. These advances pave the way for scalable hierarchical molecular storage and point toward future integrated molecular computing systems.

## Methods

### Preparation of DNA data strands and X-scaffolds

Data strands used in Figure2-Figure 4 consisted of complementary single-stranded DNA (ssDNA). To prepare the data strands, the strands were mixed equimolarly in 1×PBS buffer and annealed according to the annealing program (95℃ 5 min; 95℃-25℃, −1℃/min; 4℃ ∞) (Figure S15). Data strands in Figure 5 were prepared by two steps of PCR (Figure S16). First, we used regular PCR primers to amplify the template from oligo pools. Then, we used AP-primers containing an abasic site (AP site) to amplify the template, adding sticky ends and toehold domains to both protruding ends of the amplicons. All PCR reactions were conducted using Phusion Plus Green PCR Master Mix (Thermo Fisher Scientific, F632S) according to the manufacturer’s protocol. The X-scaffolds consisted of four partially complementary single-stranded DNA (ssDNA). To prepare the X-scaffolds, the four strands were mixed equimolarly in reaction buffer (10 mM Tris-HCl, pH 8.0) and annealed according to the annealing program (95℃ 5 min; 95℃-25℃, −1℃/min; 4℃ ∞). All ssDNA except oligo pools were synthesized by Sangon Biotech (Table S4). The Oligo pools were synthesized by Dynegene.

### Storage of DNA data strands and forming reconfigurable DNA memory

The data-enriched DNA droplets were formed by the mixture of data strands (500 nM, 1 μM, 2.5 μM, and 5 μM) and X-scaffolds (5 μM) in reaction buffer (10 mM Tris-HCl, 350 mM Na^+^, pH 8.0) at 37℃, oscillating (300 rpm) for 5-60 min (Figure S17). Data strands and X-scaffolds were both labeled with fluorescent. The mixture was centrifuged at 450 xg, 4℃ for 15 min, and the supernatant was removed to wash away excess data strands and X-scaffolds. The droplets were resuspended using the reaction buffer. After washing twice, the data-enriched DNA droplets were characterized by the laser scanning confocal microscope.

### Confocal imaging

Fluorescence data were acquired using the Zeiss confocal laser scanning microscope 800 (LSM 800) with a 63× oil-immersion objective at 512 × 512 pixels resolution. FAM was imaged with the 488 nm laser, Cy3 with the 561 nm laser, and Cy5 with the 640 nm laser. A 20 μL solution of data-enriched droplets was pipetted onto the 20 mm confocal dishes and observed at 37℃. Confocal images and fluorescence intensity profiles were analyzed using ZEN blue software.

### Analysis of the payload ratio (data/droplet)

The temporal changes of the payload ratio were analyzed from the normalized fluorescence values. First, we acquired fluorescence values from the same area inside and outside the data-enriched droplets in confocal images at 15, 30, and 60 min. To avoid the influence of background fluorescence, we subtract the outside value from the inside value. Then, we calculated the ratio of data strands to X-scaffolds and normalized the maximum to 1 and the minimum to 0.

### Fluorescence Recovery After Photobleaching (FRAP)

FRAP experiments were performed on a Zeiss LSM 800 using a 63× oil-immersion objective at 512 × 512 pixels resolution. After a 15-minute incubation, the data-enriched droplets were pipetted onto 20 mm confocal dishes. They were photobleached at 70% laser intensity at 640 nm (for Cy5-labeled data strands), and recovery was recorded every 10 seconds. Raw data measurements, background subtraction, and data normalization were conducted according to the reference^51^ to quantify droplet fluidity. We used the FRAP module of ZEN blue software to calculate the half-time of recovery (t_1/2_)^52^.

### Amplification and characterization of target DNA data strands

PCR directly amplified the data-enriched droplet samples. In a typical PCR protocol, 10 μL of 2× Phusion Plus Green PCR Master Mix (Thermo Fisher Scientific, F632S), 1 μL of sample (∼ 1 ng), 1 μL of forward primers (10 μM), 1 μL of reverse primers (10 μM), and the appropriate volume of nuclease-free water were mixed to a final volume of 20 μL. Thermal cycling started with a 30 s initial denaturation step at 98℃, followed by 30 cycles of 10 seconds at 98℃ for denaturing and 30 s at 72℃ for annealing and extension. The PCR samples were characterized by 2% agarose gel in an ice bath. After purification using the Cycle Pure Kit (Omega BioTek), the PCR products were characterized by bi-directional Sanger sequencing (Azenta).

### The encoding and decoding process

We encoded two types of files: TXT and PNG. Due to the differing file formats, each used a distinct encoding protocol. The 135 B TXT files were converted into binary strings and subsequently encoded into DNA sequences utilizing the binary-to-quaternary mapping strategy. Each DNA data strand measures approximately 120-130 nucleotides. For the 8.47 KB PNG image (350 × 341 pixels), each pixel’s RGB triplet was initially translated into a binary string and then encoded into nucleotide sequences based on tailored mapping rules. The DNA sequences obtained were then segmented into short fragments of approximately 100 nucleotides each. Each DNA data strand contains an ID domain, a payload domain, and an index domain for positional identification and data reconstruction. During the decoding process, valid reads were identified using unique primer sequences and grouped based on their index tags. The reverse mapping algorithm was then employed to convert nucleotide sequences back into binary data, thereby restoring the original files.

### Illumina sequencing

The target data strands were amplified from data-enriched droplets using primers designed to recognize the unique ID domain specifically. All amplification reactions used Phusion Plus Green PCR Master Mix (Thermo Fisher Scientific, F632S) according to the manufacturer’s protocol.

Amplicons were purified using the Cycle Pure Kit (Omega BioTek) and sent to Genefund Biotech (Shanghai, China) for library preparation and Illumina sequencing (NovaSeq X Plus).

## Data and materials availability

All data needed to evaluate the conclusions in the paper are present in the paper and/or the Supplementary Materials.

## Acknowledgements

This work was supported by National Natural Science Foundation of China grants (32571596 and 22377076), Shanghai Science and Technology Committee (25HC2810200), Shanghai Pilot Program for Basic Research and State Key Laboratory of Synergistic Chem-Bio Synthesis, Shanghai Jiao Tong University. The language of this manuscript was polished from the original text by DeepSeek.

## Author Contributions

Honglu Z. and Huan Z. conceived the project. Honglu Z. and J.Y. designed the experiments. J.Y. carried out the experiments, collected and analyzed the data. A.L. contributed to analysing the data. J.Y., Huan Z., Honglu Z. and C.F. wrote the manuscript. All authors reviewed and revised the manuscript.

## Competing interests

The authors declare that they have no competing interests.

## Notes

### Competing Interest Statement

The authors have declared no competing interest.

