## Supplementary Information for "A reconfigurable DNA memory architecture for hierarchical data management via programmable phase transitions"

**TABLE OF CONTENTS**

| Item | Pages |
|---|---|
| Supplementary Note 1 | 3 |
| Supplementary Note 2 | 3-4 |
| Supplementary Methods | 4-8 |
| Supplementary Figure S1 | 9 |
| Supplementary Figure S2 | 10 |
| Supplementary Figure S3 | 11 |
| Supplementary Figure S4 | 12 |
| Supplementary Figure S5 | 13 |
| Supplementary Figure S6 | 14 |
| Supplementary Figure S7 | 15 |
| Supplementary Figure S8 | 16 |
| Supplementary Figure S9 | 17 |
| Supplementary Figure S10 | 17 |
| Supplementary Figure S11 | 18 |
| Supplementary Figure S12 | 18 |
| Supplementary Figure S13 | 19 |
| Supplementary Figure S14 | 19 |
| Supplementary Figure S15 | 20 |
| Supplementary Figure S16 | 20 |
| Supplementary Figure S17 | 21 |
| Supplementary Figure S18 | 21 |
| Supplementary Figure S19 | 22 |
| Supplementary Table S1 | 22-23 |
| Supplementary Table S2 | 23 |
| Supplementary Table S3 | 23-24 |
| Supplementary Table S4 | 24-28 |

**Supplementary Notes**

**Supplementary Note 1. File encoding rules and workflow**

The TXT file was encoded into DNA sequences using binary-to-quaternary mapping: 00 to A, 01 to T, 10 to C, and 11 to G. Each DNA data strand is approximately 120-130 nucleotides. For the PNG image (350 × 341 pixels), the process begins with mapping each pixel's RGB triplet into a 2-bit binary string according to established conventions, resulting in a 700 × 341 binary matrix. Subsequently, each row's binary list is divided into 8-bit segments; note that the final segment comprises 4 bits. The initial 87 of these 8-bit segments are then split and converted into five nucleotide bases following Rule 2. The 88th 4-bit segment is divided into two 2-bit parts and then converted into two decimal numbers, which are then mapped to two single nucleotide bases following Rule 1. To reduce sequencing errors in long single-strand sequences, each row of bases (approximately 437 nucleotides long) is divided into four groups of 90 nucleotides, with the last group containing 77 nucleotides. Each data block is assigned an index based on its row and column positions. This decimal index is converted into a 16-bit binary code, which is then mapped to ten nucleotide bases according to Rule 1 for later indexing. Finally, each data block is given a prefix with its index label, creating an information segment of either 100 nucleotides or 90 nucleotides.

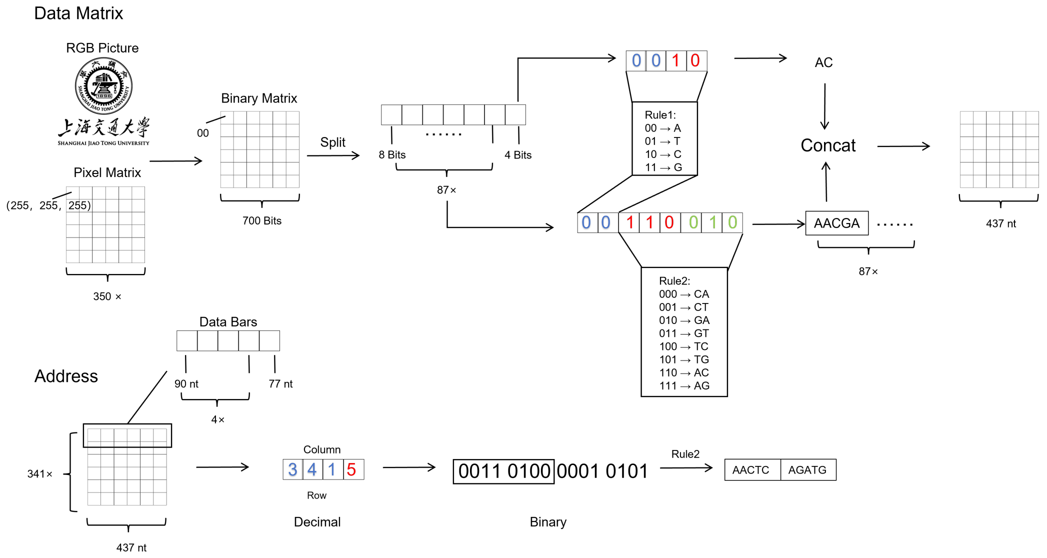

**Supplementary Note 2. File decoding rules and workflow**

Following Illumina sequencing, we decode the DNA sequences to reconstruct the original image information. The process comprises six stages: sequence reading, index parsing, matrix reconstruction, null value correction, binary restoration, and image reconstruction. We extract only the sequence content from each DNA sequence in the sequencing file. The first 10 bases function as index labels, containing row and column information encoded during the process. These are decoded using the inverse of Rule 2 to generate a 16-bit binary sequence, which is then divided into 4-bit groups and converted to decimal. The first three groups represent the row numbers (hundreds, tens, units), while the fourth specifies the column number. Based on this index, the primary segment of the DNA sequence is placed at the corresponding location in the matrix. For null or anomalous values, a "G" base sequence is used as a filler. Post-repair, each matrix row corresponds to a row of the original image. Every 5-base group is decoded into an 8-bit binary block according to Rules 1 and 2, with the final 2-base segment decoded to recover an additional 4 bits. The concatenation of all binary blocks results in a binary row sequence of length 700. Ultimately, the original image is reconstructed according to the specified color mapping.

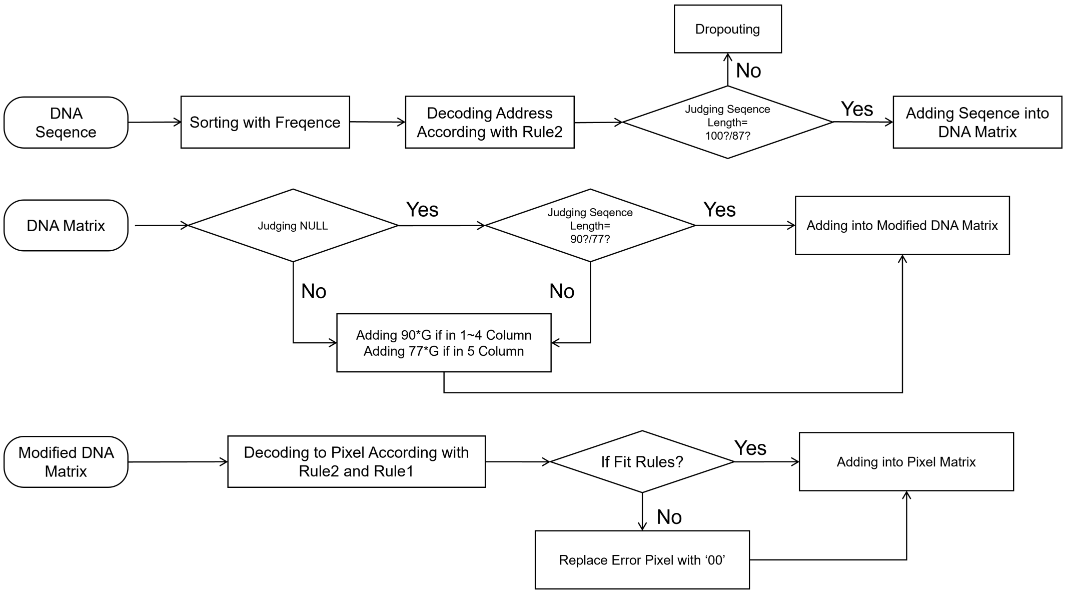

**Supplementary Methods**

**Concentration and data density in reconfigurable DNA Memory**

We establish the standard curves by measuring the fluorescence intensities of the data strands with known concentrations (Figure S18). To evaluate the concentration of data strands within droplets, we co-incubated the data strands (500 nM, 1 µM, 2.5 µM, and 5 µM) with X-scaffolds (5 µM) at

37°C, followed by two washing steps. The concentration of the data strands in the droplets was then quantified using standard calibration curves. In Figure 3b, the maximum value was normalized to 100%. For the determined length of data strands (e.g., 100 bp), utilizing the measured molar concentration ($C_n$), calculating the total number of bases ($N_{base}$) within a known volume (V) sample:

$$N_{base} = C_n \times V \times N_A \times 100 \quad (1)$$

Avogadro's constant, denoted as $N_A$, is defined as the number of particles in one mole of a substance, and its value is approximately $6.02 \times 10^{23}$ $mol^{-1}$.

In theory, each base encodes 2 bits of information, and 8 bits represent 1 byte. Thus, 4 bases represent 1 byte. This calculation enables us to ascertain the data capacity stored on the DNA strands, which we subsequently divide by the weight of the X-scaffold ($W_{scaffold}$) to determine the storage density as follows:

$$\text{Density} = \frac{N_{base}}{W_{scaffold} \times 4} \quad (2)$$

Based on our design principle, two molecules of X-scaffolds can store one molecule of data strands. Therefore, we hypothesize that 5 μM X-scaffolds can store 100 bp data strands up to 2.5 μM, and the theoretical density boundary of our system is approximately $14 \times 10^{10}$ GB/g.

**Programmable In-Memory Operations**

The programmable in-memory operations were based on toehold-mediated strand displacement reactions, including selective erasure, rewriting, replacement, and repeatable erase-rewrite cycles. For selective erasure, 10× trigger strands were added to data-enriched droplets at 37°C, with oscillation (300 rpm) for 5-20 min, to release the target data strands. This process involved recognizing the unique toehold domains in the target data strands and competitively replacing the sticky ends between data strands and droplets. For rewriting and replacement, the post-erasure samples were washed twice and then loaded with either the original or updated data strands (500 nM, 1 μM, 2.5 μM, and 5 μM). They were subsequently incubated at 37°C with oscillation (300 rpm) for 15 min to facilitate data rewriting and replacement. For repeatable erasure-rewrite cycles, the samples were subjected to the conditions mentioned above, undergoing a series of selective erasure, washing, rewriting, and washing cycles. After each erasure and rewriting, the samples were washed, and their fluorescence intensities measured. The concentration of data strands in the samples after each operation was calculated based on the established fluorescence intensity-concentration standard curve.

**Quantitative analysis of retention data**

Following in-memory operations, the retention data in droplets were quantified by qPCR. All qPCR analyses were conducted using Phusion Plus Green PCR Master Mix (Thermo Fisher Scientific, F632S) according to the manufacturer's protocol, using 96-well plates on the Archimed-X4 qPCR machine (RocGene). The known concentration data strands were used to create the standard curves (Figure S19) for measuring retention concentration. The retention ratio of data strands in droplets was determined by dividing the measured concentration (C) by the original concentration ($C_0$).

**Calculating editing efficiency from sequencing results**

First, read the original image's domain2 and the edited domain2 independently, converting both to RGB mode. Utilize all pixels with RGB values (0,0,0) in the original image as the target region. Subsequently, compare the pixel values at corresponding positions between the original and modified images on a pixel-by-pixel basis, and count the number of black pixels that have changed. The editing efficiency was determined following the equation:

$$\text{Editing Efficiency} = 1 - \frac{N_{\text{changed}}}{N_{\text{black}}} \times 100\% \tag{3}$$

Here, $N_{black}$ represents the total number of black pixels, and $N_{changed}$ means the number of changed pixels.

**Calculating the correct decoding rate from sequencing results**

First, read the original and restored images separately, converting both to RGB mode. Utilizing all pixels in the original image as the reference area, perform a pixel-by-pixel comparison between the original and restored images at corresponding positions, and identify regions where the pixel values are consistent. The correct decoding efficiency was determined following the equation:

$$\text{Decoding Efficiency} = \frac{N_{\text{same}}}{N_{\text{Original}}} \times 100\% \tag{4}$$

Here, $N_{Original}$ represents the total number of pixels in the original image, and $N_{same}$ represents the number of pixels in the restored images that match the original.

**Reversible encapsulated by the tetrahedral DNA framework**

After in-memory operations, the hot data need transition into a stable, archival cold data state for long-term preservation. We design the tetrahedral DNA framework (TDF) to reversibly encapsulate the data-enriched droplets. During encapsulation, we added bridge strands and TDF to data-enriched droplets, ensuring a 10:2:1 molar ratio of X-scaffolds, bridge strands, and TDF in the system. After incubation at 37°C for 15 minutes, core-shell nanostructures formed. To remove the TDF's shell, we add the excess release strand to the decapsulated droplet sample and incubate at 37°C for 15 minutes, maintaining a 10:1 molar ratio of release strands to TDF in the reaction.

**Digestion by enzymes**

To test the protective efficiency of TDF's shell, samples (data strands in free solution, decapsulated droplets, and encapsulated droplets) containing 500 nM data strands were treated with 50 U/mL EcoRI (NEB) or 10 U/mL DNase I (Sangon Biotech) in a 30 μL reaction volume. The reactions were incubated at 37°C for 0, 5, 10, 20, and 30 min. The data strands contain two EcoRI recognition sites. After digestion, the data fragment was cut between the ID domains at both ends, preventing amplification of the target information by ID domain recognition. After stopping the digestion reactions by heating, the retention data were measured by qPCR.

**Accelerated ageing experiment and analysis**

To test long-term stability, the samples (data strands in free solution, decapsulated droplets, and encapsulated droplets) were placed in a thermocycler (Thermo Fisher Scientific) at 65°C for 0, 8, 16, 24, and 48 h. After treatment, the retention data were measured by qPCR, and the decay rate (k) was calculated using the following equation:

$$\ln\left(\frac{[C]}{[C_0]}\right) = -kt \tag{5}$$

Here, $C_0$ is the initial concentration of data strands at the 0-hour time point, and C is the concentration measured at each subsequent time point. Then perform linear regression on the data points using the equation above; the slope of the resulting linear model represents k. Subsequently, we calculated the half-life using the following equation:

$$t_{\frac{1}{2}} = \frac{\ln(2)}{k} \quad (6)$$

Based on the Arrhenius extrapolation models from Grass et al.[1], we estimate that the storage half-life of data in our encapsulated droplets system is approximately 4,000 years at 4°C and approximately 1.4 million years at -18°C.

**Cryopreservation stress testing**

The 30 μL data-enriched droplet samples were frozen at -20°C, -80°C, and -196°C (liquid nitrogen) for 12h, respectively. Subsequently, the frozen samples were completely thawed at 37°C, oscillating (300 rpm) for 15 min. We observed the morphology and population of thawed droplets using confocal microscopy and compared them to the control group. All data were analyzed with ImageJ.

### Supplementary Figures

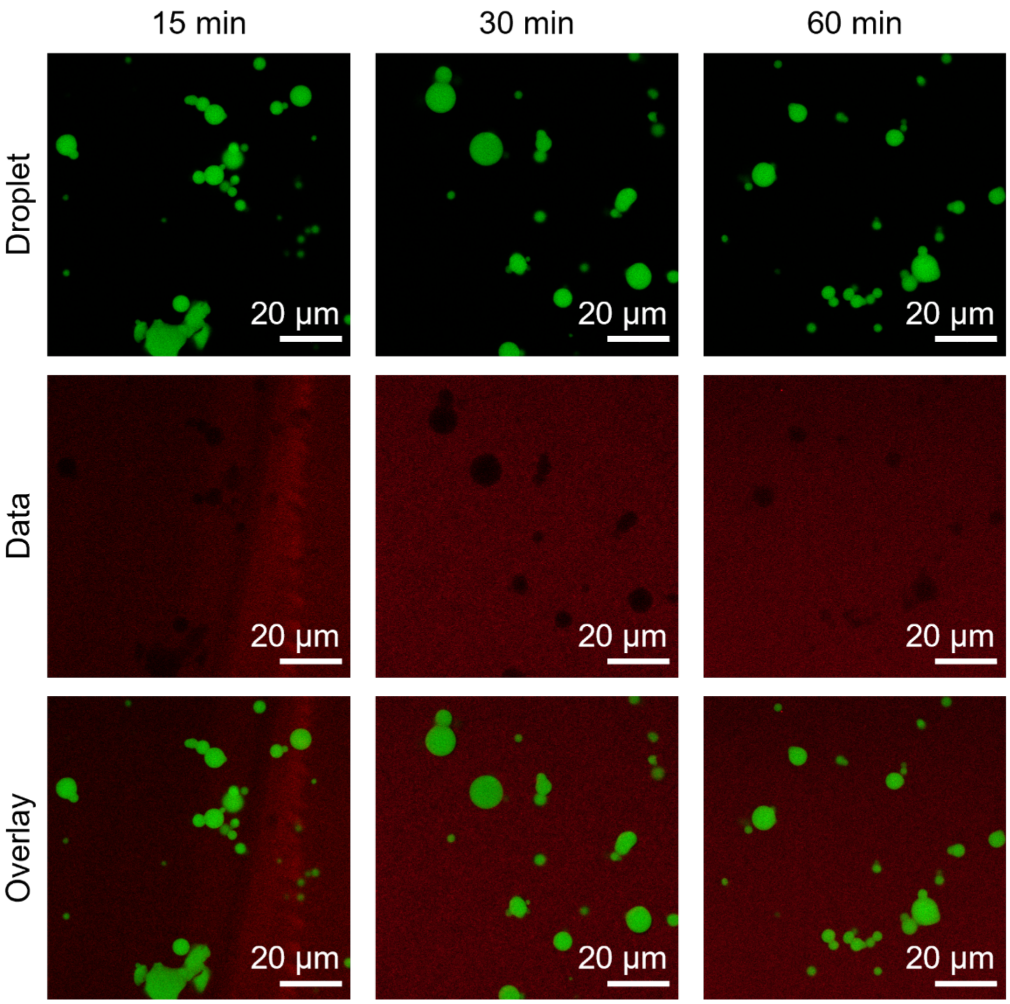

**Figure S1. Temporal fluorescence images of data strands (50 bp dsDNA) stored in the P-4 system.**

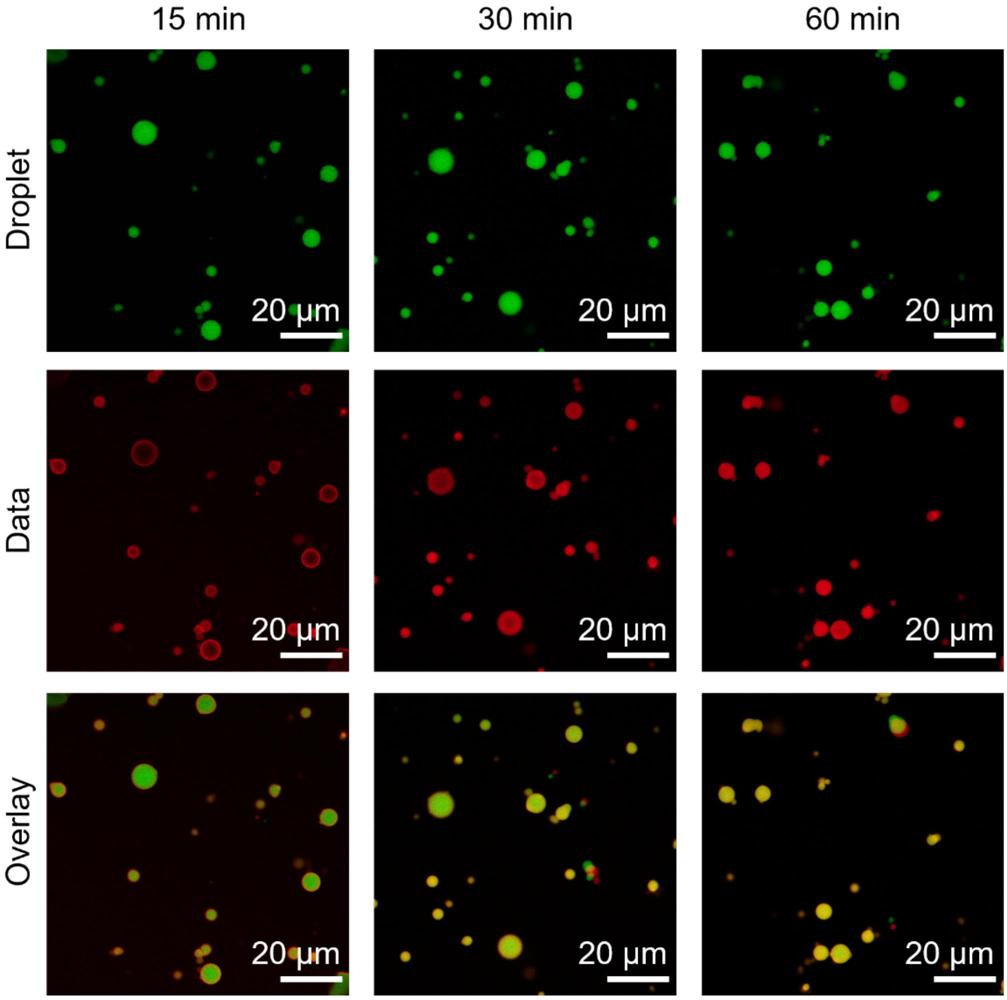

**Figure S2. Temporal fluorescence images of data strands (50 bp dsDNA) stored in the P-6 system.**

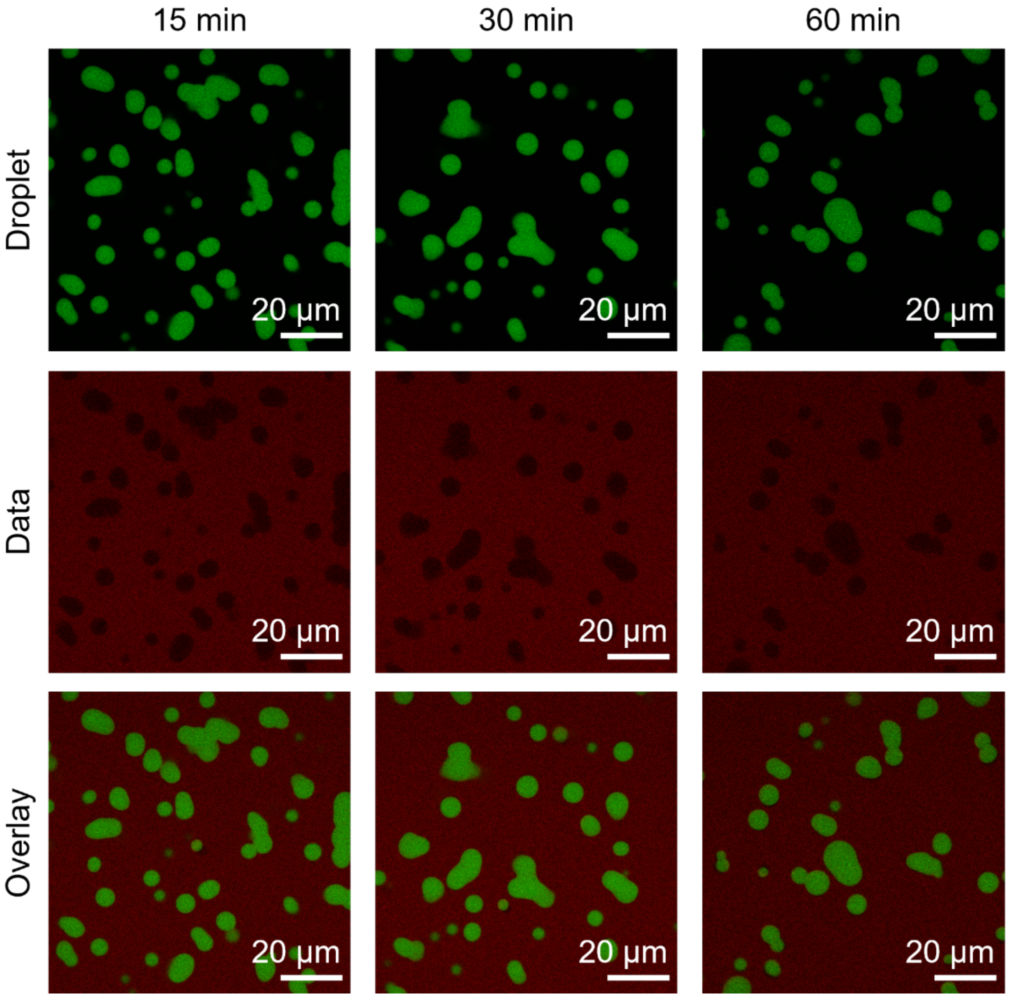

**Figure S3. Temporal fluorescence images of data strands (50 bp dsDNA) stored in the NP-4 system.**

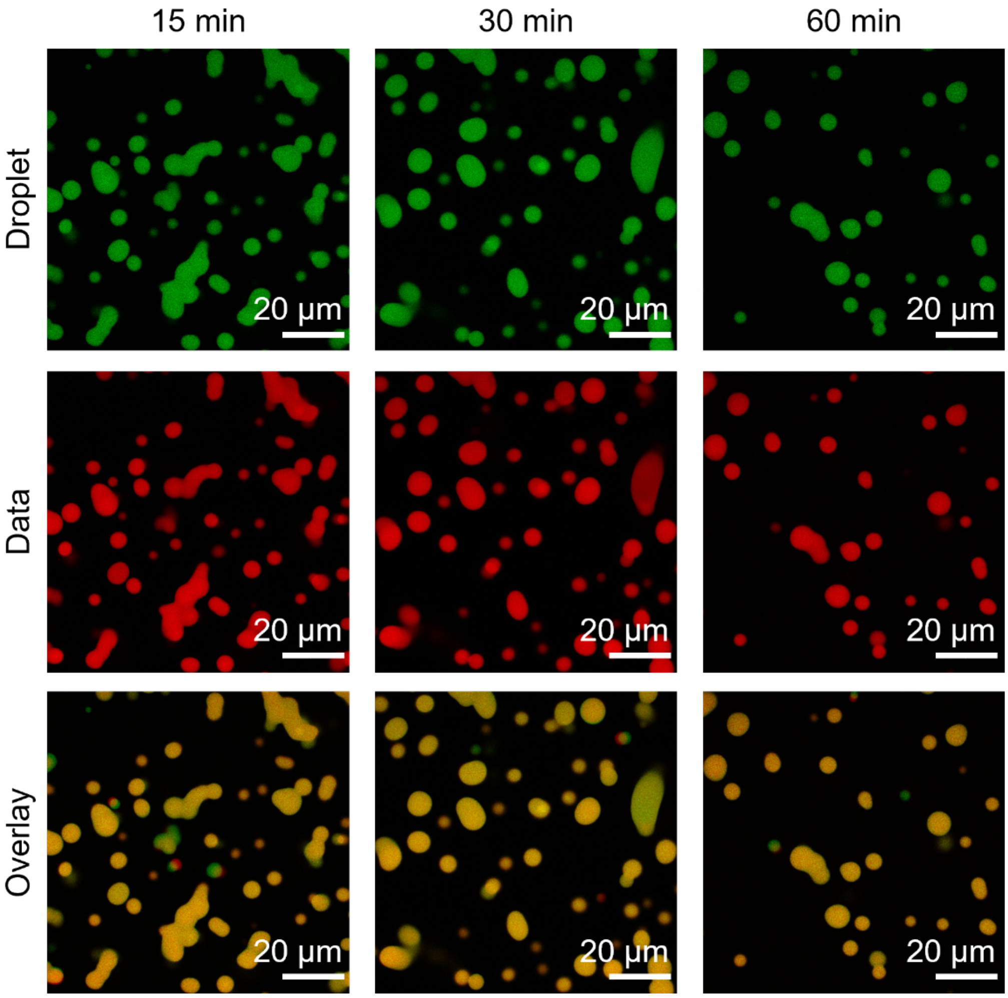

**Figure S4. Temporal fluorescence images of data strands (50 bp dsDNA) stored in the NP-6 system.**

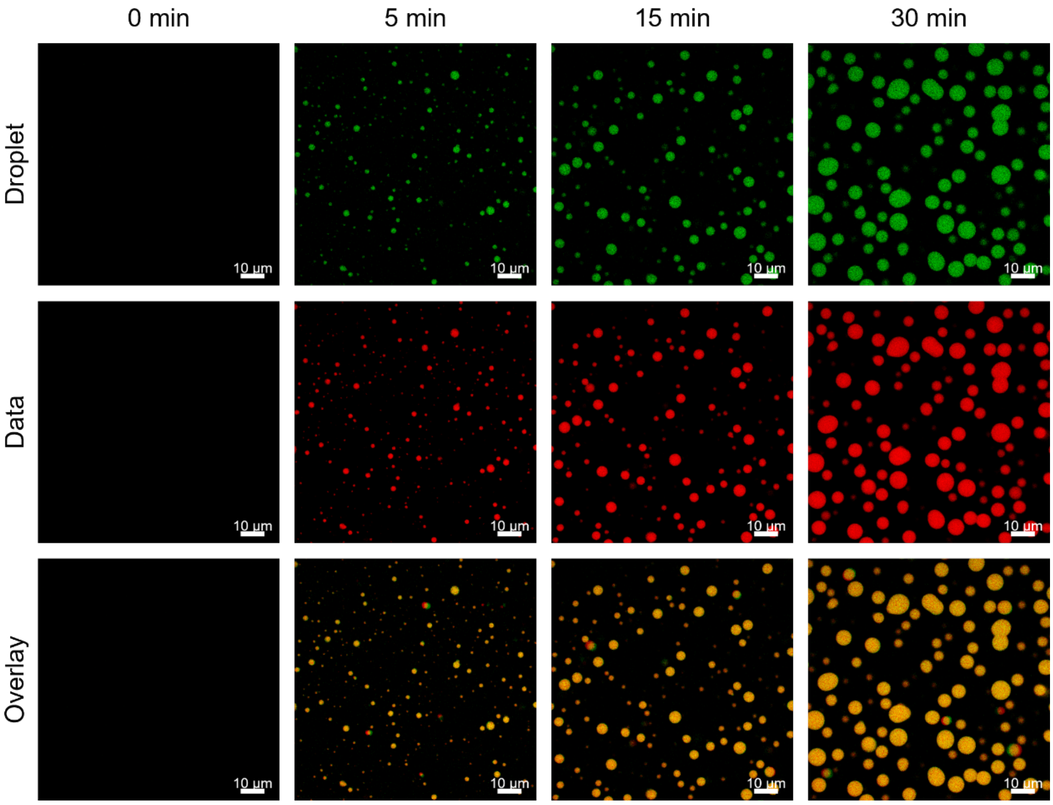

**Figure S5. Temporal fluorescence images of data strands (100 bp dsDNA) stored in the NP-6 system.**

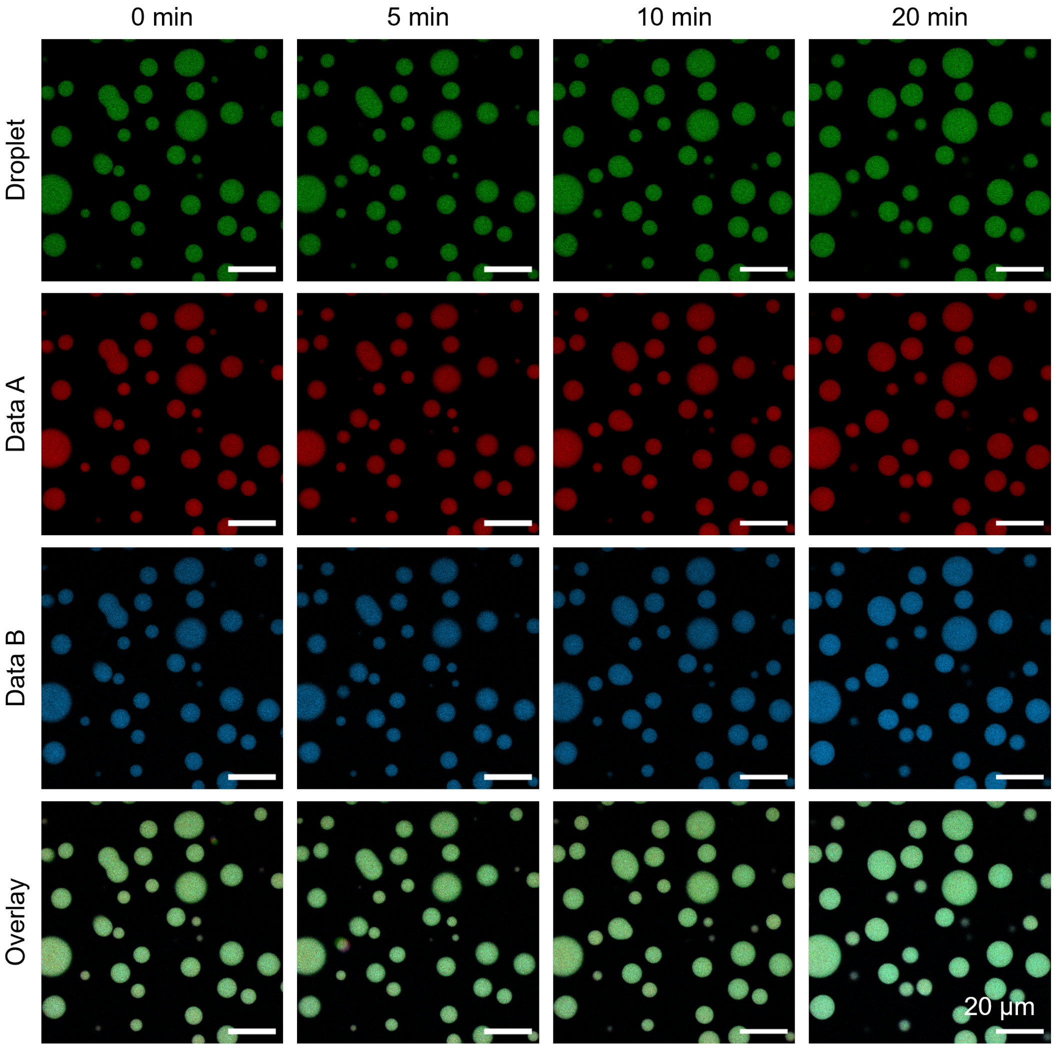

**Figure S6. Temporal fluorescence images of the NOT gate (Control) during selective erasure.**

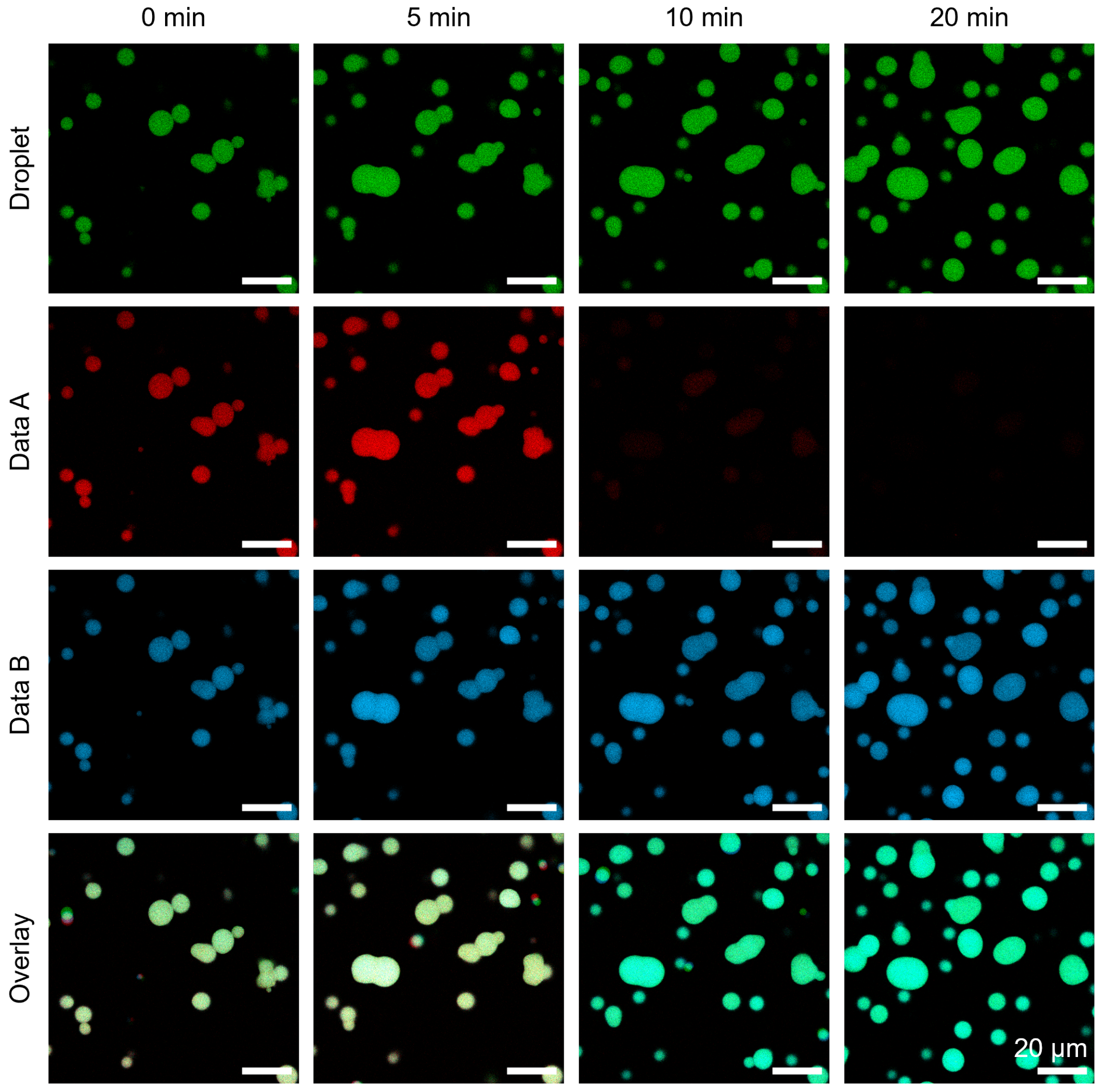

**Figure S7. Temporal fluorescence images of the OR gate (Delete A) during selective erasure.**

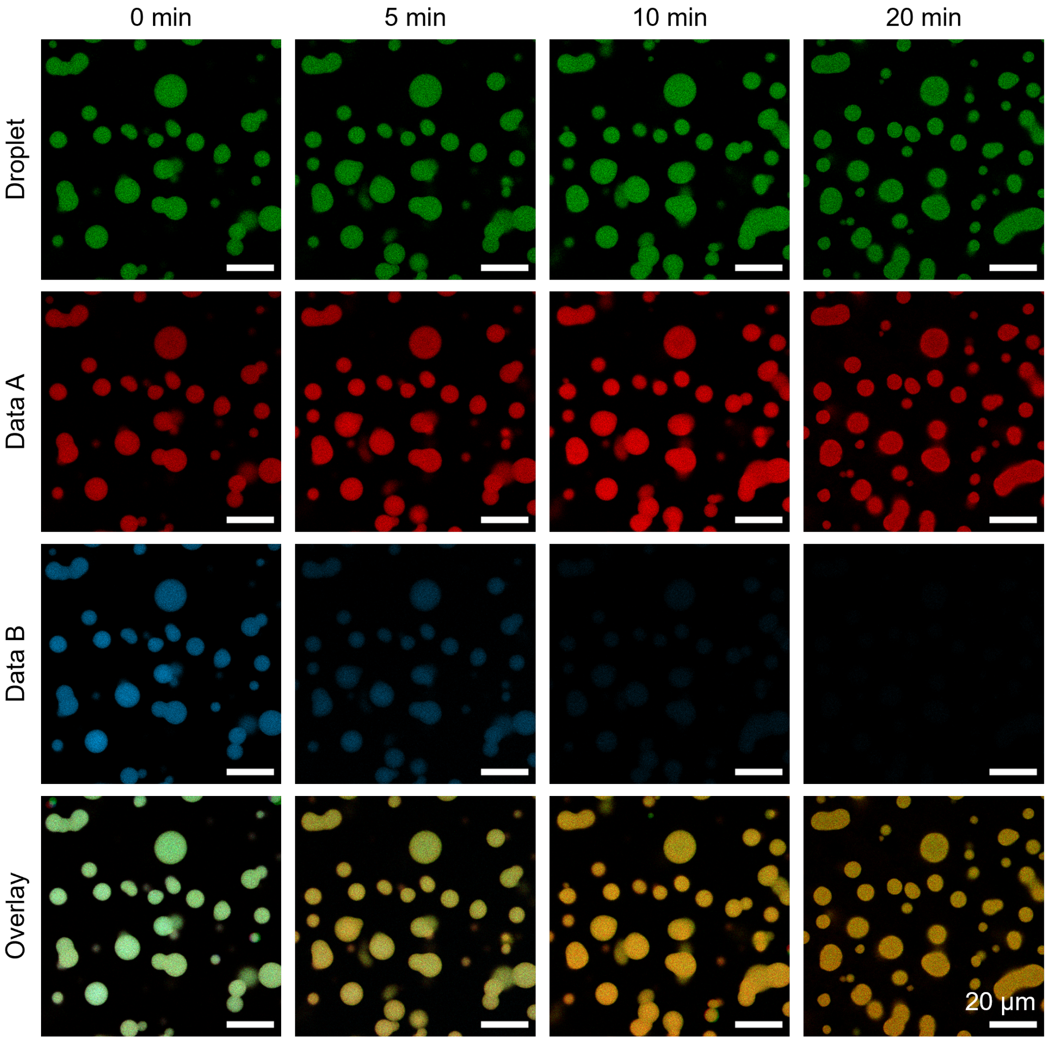

**Figure S8. Temporal fluorescence images of the OR gate (Delete B) during selective erasure.**

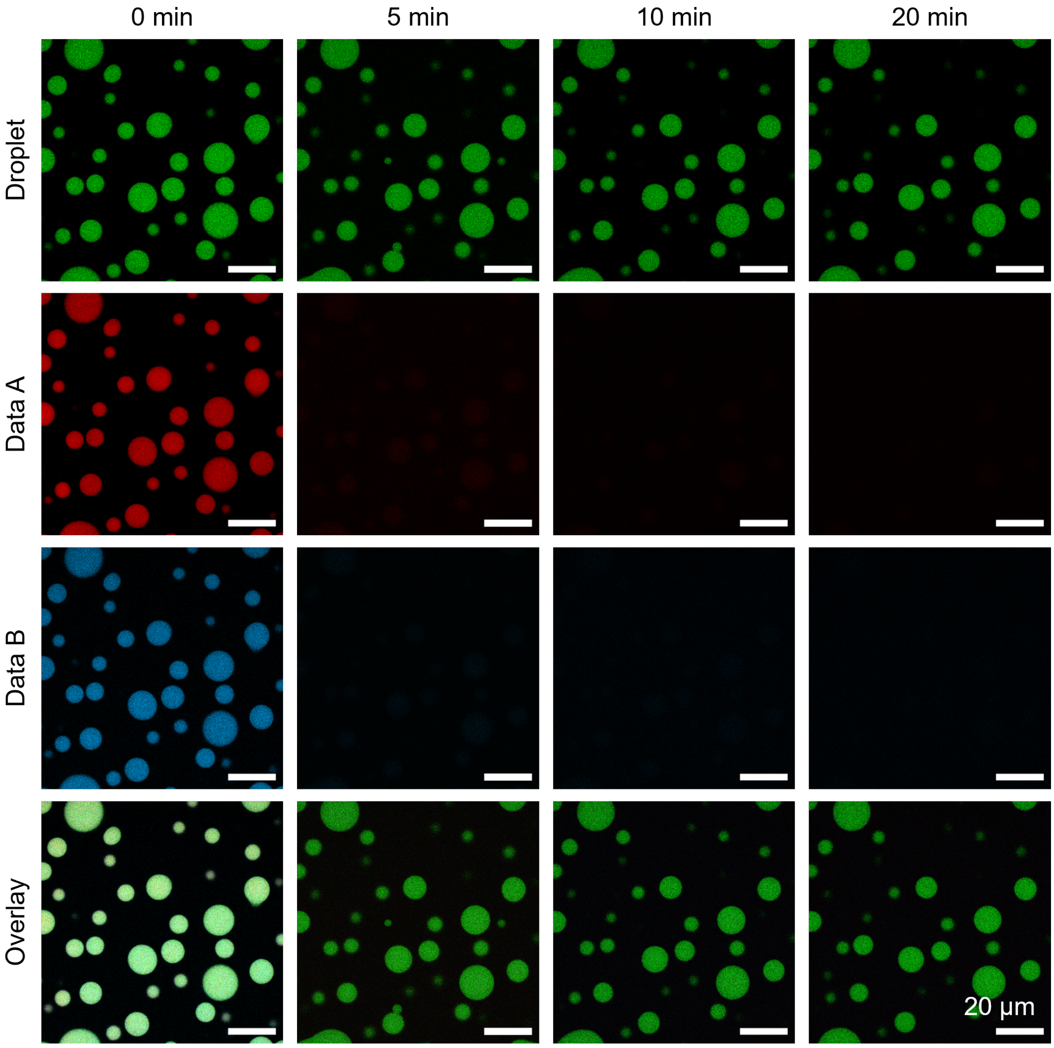

**Figure S9. Temporal fluorescence images of the AND gate (Delete A & B) during selective erasure.**

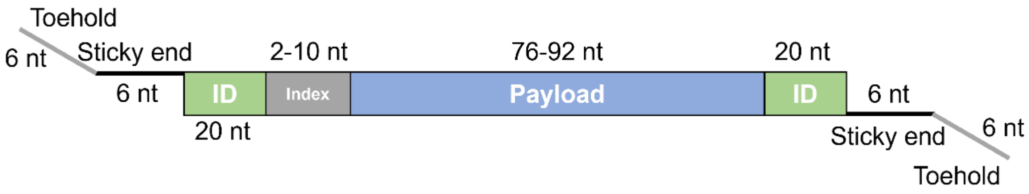

**Figure S10. Composition of DNA data strands for TXT and PNG files.**

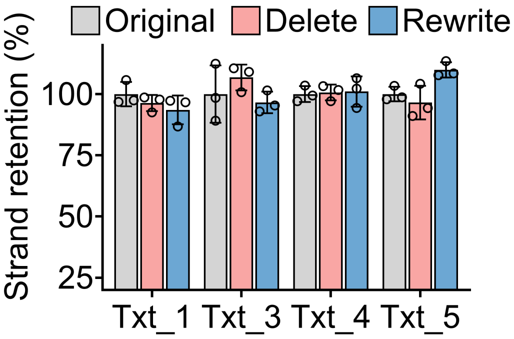

**Figure S11. Retention ratios of non-target strands (Txt_2a, Txt_6a) following selective deletion and rewriting operations.** The non-target strands remained unaffected. Error bars represent SD (n = 3).

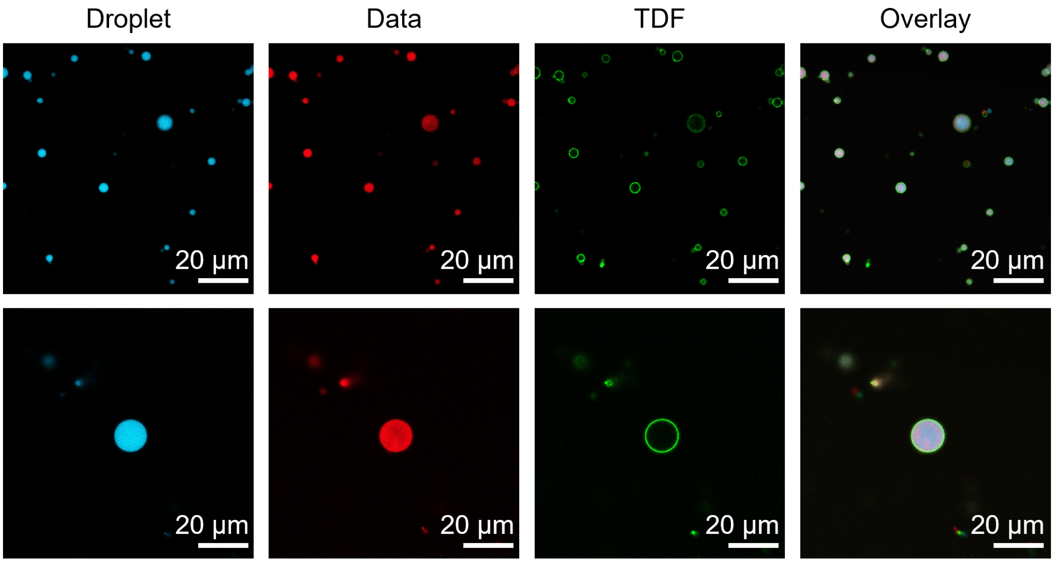

**Figure S12. The fluorescence images of the encapsulated droplets.**

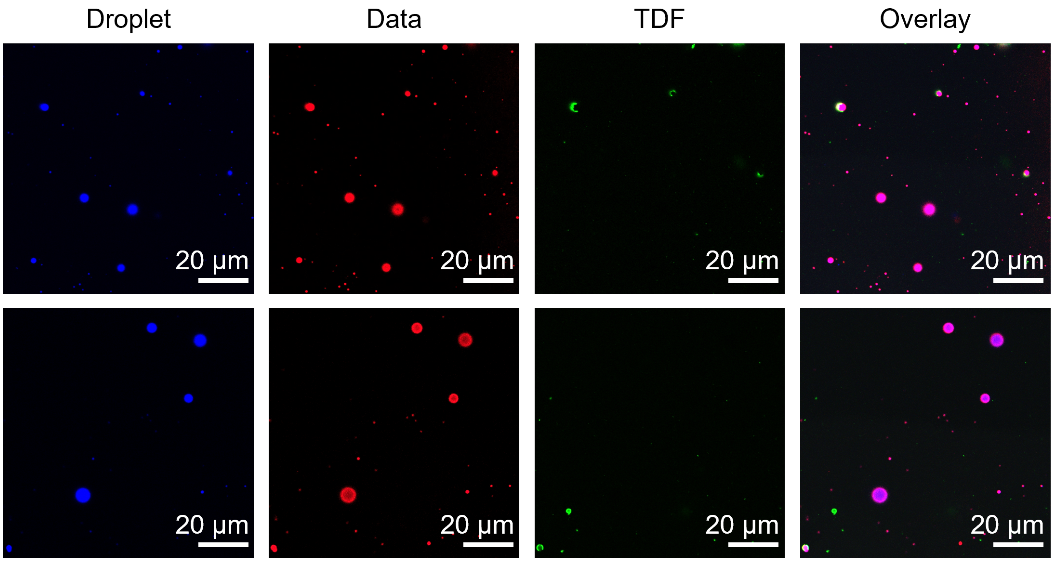

**Figure S13. The fluorescence images of the decapsulated droplets.**

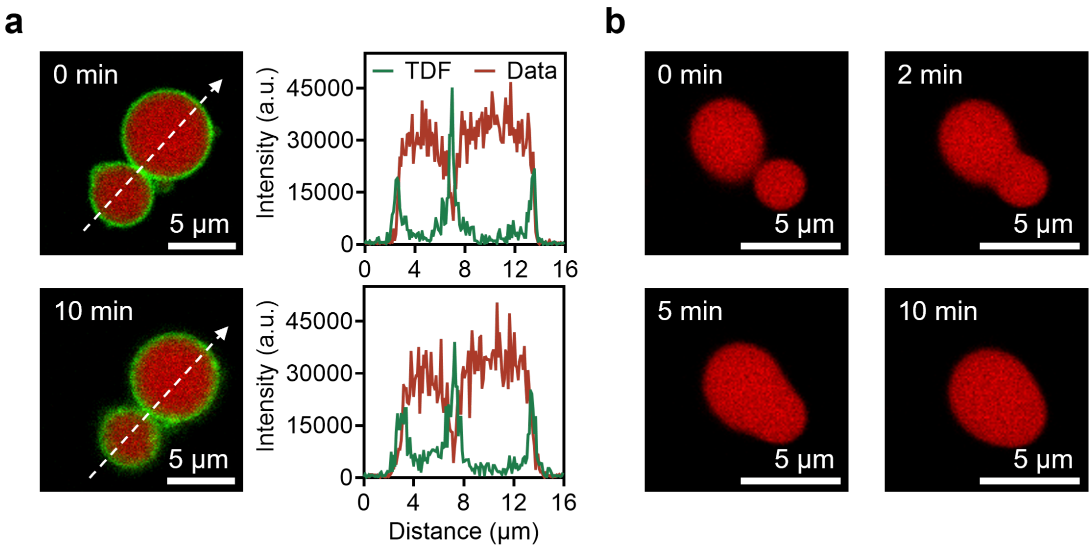

**Figure S14. TDF encapsulation prevents droplet fusion and maintains colloidal stability. a,** Temporal fluorescence images (left) and corresponding intensity profiles (right) of encapsulated droplets. **b**, Temporal fluorescence images of decapsulated droplets.

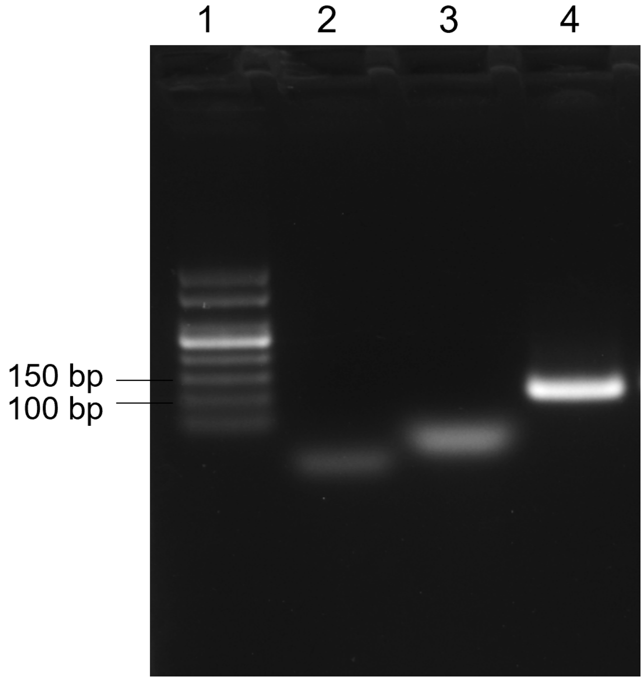

**Figure S15. 2% agarose electrophoresis analysis of the annealed data strands.** Line 1: 50-500 bp marker; line 2: primer-F; line 3: primer-R; line 4: data strand.

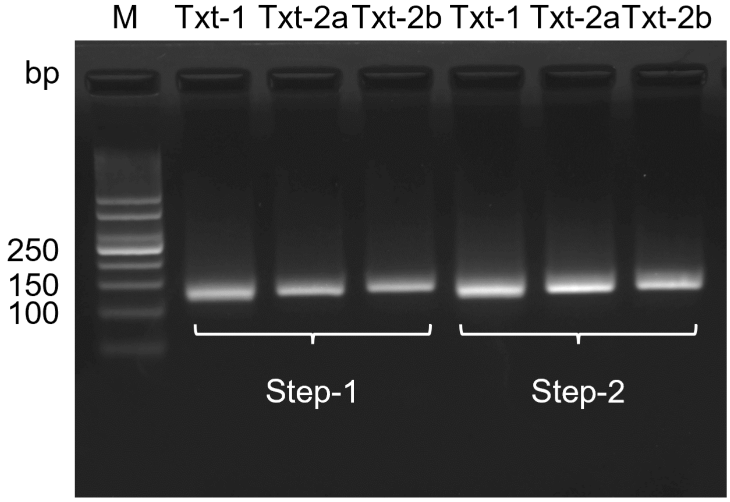

**Figure S16. 2% agarose electrophoresis analysis of the data strands prepared by two-step PCR.**

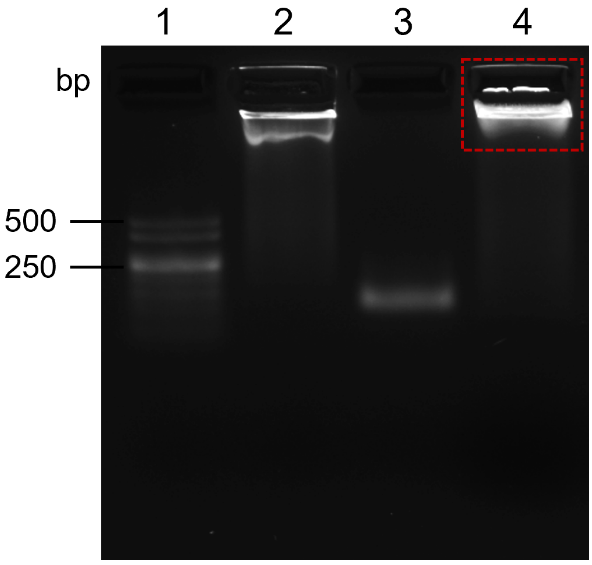

**Figure S17. 2% agarose electrophoresis analysis of the data storage.** Line 1: 50-500 bp marker; line 2: X-scaffolds; line 3: data strands; line 4: X-scaffolds + data strands.

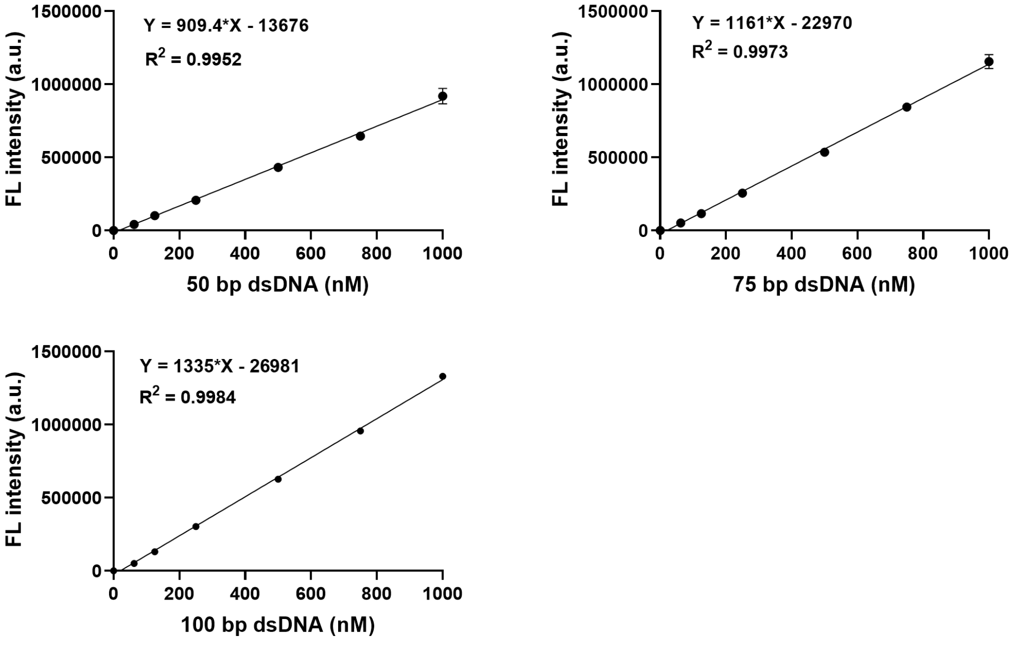

**Figure S18. Standard curves of DNA concentrations versus fluorescence intensity for different lengths (50, 75, 100 bp).**

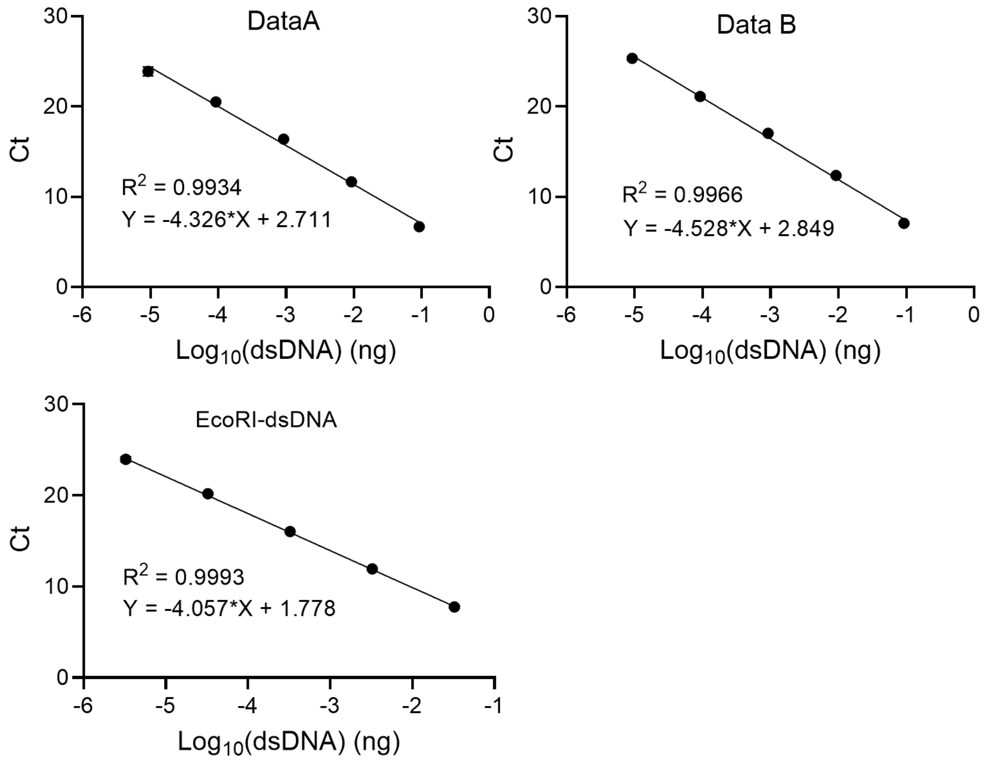

**Figure S19. Standard curves of $Log_{10}$ (data mass) versus cycle threshold (Ct) for distinct dsDNA data strands.**

### Supplementary Tables

**Table S1. Comparison of different media for DNA data storage**

| Storage media | Data density | Function |
|---|---|---|
| Theoretical boundary of the DNA molecule[2] | $4.55\times10^{11}$ GB/g | Storage |
| Theoretical boundary of this work | $1.4\times10^{11}$ GB/g | Storage and In-memory editing |
| **This work** | $7\times10^{10}$ GB/g | Storage and In-memory editing |
| SSD[3] | 60 GB/g | Storage and In-memory editing |
| $SiO_2$[4] | $6.86\times10^{6}$ GB/g | Storage |
| TRFG hydrogels[5] | $7\times10^{9}$ GB/g | Storage |
| Multilayer Magnetic Nanoparticles[3] | $8\times10^{9}$ GB/g | Storage |

| | | |
|---|---|---|
| Photonic microspheres[6] | $2.43\times10^{10}$ GB/g | Storage and access |
| Core–shell DNA condensates[7] | $9.23\times10^{2}$ GB/g* | Storage and encryption |

*To calculate the data density of core–shell DNA condensates[7], we assume the condensate density is comparable to water (1 g/cm). Based on the estimated upper-bound volumetric information density of $7.93\times10^{9}$ bits/mm$^3$, the corresponding mass-normalized storage density is approximately $9.23\times10^{2}$ GB/g.

**Table S2. The decay rate (k) and half-life ($t_{1/2}$) of data strands at 65°C**

| Sample | Decay Rate k ($s^{-1}$) at 65°C | Half-Life $t_{1/2}$ (years) |
|---|---|---|
| Free 100 bp dsDNA | $4.69\times10^{-6}$ | 0.00469 |
| Droplet | $1.6\times10^{-6}$ | 0.0137 |
| Blocked droplet | $1.07\times10^{-6}$ | 0.0205 |

**Table S3. Comparison of performance metrics for different dynamic DNA storage systems**

| **Architecture** | **Editing location** | **Editing granularity** | **Protection mechanism** | **Hot-cold memory hierarchy** |
|---|---|---|---|---|
| **This work** | In-memory (condensate-based) | Fine (bit-level) | DNA framework encapsulation | Yes (reversible) |
| ss-dsDNA in solution[8] | Off-memory (bulk solution) | Medium (strand-level) | None | No |
| Plasmid in solution[9] | In-memory (plasmid) | Medium (logic-level) | None | No |
| Liquid metal[10] | Off-memory (release-recapture) | Coarse (file-level) | Heating-induced phase change | No |
| Core-shell | In-memory | Medium | None | No |

| | | |
|---|---|---|
| condensates[7] | (spatially-based) | (spatial-level) |

**Table S4. Sequence of X-scaffolds, data strands, primers, and TDF**

| Name | Sequence (5’-3’) |
|---|---|
| X-1 | CGCGAGCAAACAAGTCTACGAATCAATTCCCGTTGGCT |
| X-2 | CGCGATTACAGACCCTCTCAAACCGAGCATATGTCAGA |
| X-3 | CGCGTCTGACATATGCTCGGAACGTAGACTTGTTTGCT |
| X-P4 | CGCGAGCCAACGGGAATTGAAATGAGAGGGTCTGTAAT |
| X-P6 | CGCGCGAGCCAACGGGAATTGAAATGAGAGGGTCTGTAAT |
| X-NP4 | ACCTAGCCAACGGGAATTGAAATGAGAGGGTCTGTAAT |
| X-NP6 | TGACCTAGCCAACGGGAATTGAAATGAGAGGGTCTGTAAT |
| P50-F | CGCGCGAGCACTCTTCGAAATAGCTTATTCACGTCGTGCCACATTACGAACGCTTC |
| P50-R | CGCGCGGAAGCGTTCGTAATGTGGCACGACGTGAATAAGCTATTTCGAAGAGTGCT |
| NP50-F | AGGTCAAGCACTCTTCGAAATAGCTTATTCACGTCGTGCCACATTACGAACGCTTC |
| NP50-R | AGGTCAGAAGCGTTCGTAATGTGGCACGACGTGAATAAGCTATTTCGAAGAGTGCT |
| NP75-F | AGGTCACTTCGCACAATTAGTCCTTGATCTTACAATATATAGCTACAATCGTATCATACAGTGTATCGGTGACTCACTGTC |
| NP75-R | AGGTCAGACAGTGAGTCACCGATACACTGTATGATACGATTGTAGCTATATATTGTAAGATCAAGGACTAATTGTGCGAAG |
| NP100-F | AGGTCAAACAAGACATGGAATAGCGTGCCACAGCTTATGCGGGATGAGGAATATAAGTGGCTGATGACCCCCCTTTCTGGAGCTCTTCAGCACCGCAGCGAGAACA |

| | |
|---|---|
| NP100-R | AGGTCATGTTCTCGCTGCGGTGCTGAAGAGCTCCAGAAAGGGGGGTCATCAGCCACTTATATTCCTCATCCCGCATAAGCTGTGGCACGCTATTCCATGTCTTGTT |
| Data A-F | GACTAGAGGTCACTACTGTCTGCGTTGAGACTTATATAGCTAATACAATATATCATTGTATCATACAATATTTCTATCCTTGTATCCTTCGCTCTGAGAACAATACGGACAATCGG |
| Data A-R | GACTAGAGGTCACCGATTGTCCGTATTGTTCTCAGAGCGAAGGATACAAGGATAGAAATATTGTATGATACAATGATATATTGTATTAGCTATATAAGTCTCAACGCAGACAGTAG |
| Data B-F | GCTTGAAGGTCATCTCGAGTCGTGATATGAAGTATATAGCTAATACAATATATCATTGTATCATACAATTAGTGTATCGGTGACTCATTCTGTCTTCAATCCTCATAATTGCCTGC |
| Data B-R | GCTTGAAGGTCAGCAGGCAATTATGAGGATTGAAGACAGAATGAGTCACCGATACACTAATTGTATGATACAATGATATATTGTATTAGCTATATACTTCATATCACGACTCGAGA |
| EcoRI-F | CGCGCGGTCTAGAGATGACAGTGCGAATTCTATATAGCTAATTGTATCATACAATTAGTGTATCGGTGACTCATTCTGTCTTGAATTCTCGTGATAGCTCATCACC |
| EcoRI-R | CGCGCGGGTGATGAGCTATCACGAGAATTCAAGACAGAATGAGTCACCGATACACTAATTGTATGATACAATTAGCTATATAGAATTCGCACTGTCATCTCTAGAC |
| Pri-Edit-F | CTACTGTCTGCGTTGAGACT |
| Pri-Edit-R | CCGATTGTCCGTATTGTTCT |
| Pri-Data A-F | CTACTGTCTGCGTTGAGACT |
| Pri-Data A-R | CCGATTGTCCGTATTGTTCT |
| Pri-Data B-F | TCTCGAGTCGTGATATGAAG |
| Pri-Data B-R | GCAGGCAATTATGAGGATTG |

| | |
|---|---|
| Pri-EcoRI-F | GTCTAGAGATGACAGTGCGA |
| Pri-EcoRI-R | GGTGATGAGCTATCACGAGA |
| Trigger A | TGACCTCTAGTC |
| Trigger B | TGACCTTCAAGC |
| TXT-1 | GCCTTATAATCGCGTCCTAGTTTATCGGACAATGAGTCTTTCTTACA<br>ATCATACAATGTGTCGGTGACTCGATCTAACAATCCTTCGCACAATC<br>ATACAAATTGGTGTGGGACTTACAAGGT |
| TXT-2a | GCCTTATAATCGCGTCCTAGTCTGTGACTCATTCCTTCGCACAATCG<br>GTCTCACAATGAGTCATTCGCTCTAACGATAATTCGCTCTAACAATC<br>ATACAATCCATCTTACTGGTGTGGGACTTACAAGGT |
| TXT-2b | GCCTTATAATCGCGTCCTAGTCTATGACTCGGTGAATCGATCTTTGT<br>AACAATCGGTCTCACAATATATAGCTAATACGATAATTCGCTCTAAC<br>AATCATACAATCCATCTTACTGGTGTGGGACTTACAAGGT |
| TXT-3 | GCCTTATAATCGCGTCCTAGTCATTGTCTCTTTCGCACAATCCTTCG<br>CACAATCATACAATGTGTCCTTCGATCTAACAATCTCTCGATCGGTG<br>TGTCTTTGACACGAAGTGGTGTGGGACTTACAAGGT |
| TXT-4 | GCCTTATAATCGCGTCCTAGTACATCGGTCGATCTAACAATCCTTCG<br>CTCTCTCCTTCGCTCCTTGTATGCTACAATCCTTCGCACAATGTATC<br>CATCTTACAATGAATATGGTGTGGGACTTACAAGGT |
| TXT-5 | GCCTTATAATCGCGTCCTAGTCATTCGATCGTACAATCGGTCTCACA<br>ATGCTTCGGTGTTTGACACAATCCATCATTCGCTCTAACGATAATTC<br>GCTCTAACAATCTTTTTGGTGTGGGACTTACAAGGT |
| TXT-6a | GCCTTATAATCGCGTCCTAGTGTATCTTTGACTCGCTCCTTGTATGCT<br>ACAATCCTTCGCACAATCATTCGCACAATCCATCGGTGTTTGACAC<br>GCTCTGGTGTGGGACTTACAAGGT |
| TXT-6b | GCCTTATAATCGCGTCCTAGTGTATCTTTGACTCGCTCCTTGTATGCT<br>ACAATCCTTCGCACAATCATACAATCGTTCGGTCGTTCTTTCGCTGT |

| | |
|---|---|
| | AACGCTCTGGTGTGGGACTTACAAGGT |
| Pri-Txt-F | GCCTTATAATCGCGTCCTAG |
| Pri-Txt-R | ACCTTGTAAGTCCCACACCA |
| Pri-Png-F | AACCTGTCTTTCCTAGCTCC |
| Pri-Png-R | GTCGTGTCATGGCTTTACCA |
| Txt-M-ASPF | TGGAAGAGGTCA/idSp/GCCTTATAATCGCGTCCTAG |
| Txt-M-ASPR | TGGAAGAGGTCA/idSp/ACCTTGTAAGTCCCACACCA |
| Txt-2-ASPF | ACTTCCAGGTCA/idSp/GCCTTATAATCGCGTCCTAG |
| Txt-2-ASPR | ACTTCCAGGTCA/idSp/ACCTTGTAAGTCCCACACCA |
| Txt-6-ASPF | CGACATAGGTCA/idSp/GCCTTATAATCGCGTCCTAG |
| Txt-6-ASPR | CGACATAGGTCA/idSp/ACCTTGTAAGTCCCACACCA |
| Png-D1-ASPF | ACAGTCAGGTCA/idSp/AACCTGTCTTTCCTAGCTCC |
| Png-D1-ASPR | ACAGTCAGGTCA/idSp/GTCGTGTCATGGCTTTACCA |
| Png-D2-ASPF | GGCTATAGGTCA/idSp/AACCTGTCTTTCCTAGCTCC |
| Png-D2-ASPR | GGCTATAGGTCA/idSp/GTCGTGTCATGGCTTTACCA |
| Trigger-Txt-M | TGACCTCTTCCA |
| Trigger-Txt-2 | TGACCTGGAAGT |
| Trigger-Txt-6 | TGACCTATGTCG |
| Trigger-Png-D1 | TGACCTGACTGT |
| Trigger-Png-D2 | TGACCTATAGCC |
| B-T20-a | ATTTATCACCCGCCATAGTAGACGTATCACCAGGCAGTTGAGACGA ACATTCCTAAGTCTGAAAAGCTCGAGC |
| B-T20-b | ACATGCGAGGGTCCAATACCGACGATTACAGCTTGCTACACGATTC AGACTTAGGAATGTTCGAAAAAAAAAA |

| | |
|---|---|
| B-T20-c | ACGGTATTGGACCCTCGCATGACTCAACTGCCTGGTGATACGAGGATGGGCATGCTCTTCCCGAAGCTCGAGC |
| B-T20-d | ACTACTATGGCGGGTGATAAAACGTGTAGCAAGCTGTAATCGACGGGAAGAGCATGCCCATCCAAGAAGAGGGTCATCAGT |
| Bridge-T20 | CGCGACTGATGACC |
| Release-T20 | ACTGATGACCCTCTTC |
